# Convergent morphology and divergent phenology: unravelling the coexistence of mimetic *Morpho* butterfly species

**DOI:** 10.1101/2020.11.26.399931

**Authors:** Camille Le Roy, Camille Roux, Elisabeth Authier, Héloïse Bastide, Vincent Debat, Violaine Llaurens

**Affiliations:** Institut de Systématique, Evolution, Biodiversité (ISYEB), Muséum National d’Histoire Naturelle, CNRS, Sorbonne Université, EPHE, Université des Antilles, CP50, 75005 Paris, France; Université Paris Descartes, Sorbonne Paris Cité, 12 rue de l’École de Médecine, 75006 Paris, France; Université Lille, CNRS, UMR 8198 - Evo-Eco-Paleo, F-59000 Lille, France; Laboratoire Evolution Génomes Comportement et Ecologie, CNRS, IRD, Université Paris-Sud, Université Paris-Saclay, Gif-sur-Yvette, France

**Keywords:** Reproductive interference, Territoriality, Mate recognition, Cryptic signalling, Escape mimicry

## Abstract

The emergence and persistence of closely-related species in sympatry is puzzling because the potential gene flow and the common local selective pressures may lead to either merging or competitive exclusion. Some species of *Morpho* butterflies occurring in sympatry display highly similar wing colour patterns. Associated with erratic flight abilities, their bright colouration may limit predator success and discourage future attacks. The evolution of similar colouration in sympatric species is thus likely under local selection by predators (i.e. escape mimicry). Such phenotypic similarity may promote interspecific territoriality and/or reproductive interference, questioning how closely-related co-mimetic species become sexually isolated and coexist in sympatry. We performed a series of field experiments using flying *Morpho* dummies placed in a natural habitat where wild males commonly patrol. Analysing the interactions of wild *Morpho* with different dummies, we show that similarity in wing colour pattern leads to interspecific territoriality and courtship among sympatric species. Using genomic data, we then showed that sympatric *Morpho* species are surprisingly strictly isolated despite their close relatedness and the observed heterospecific interactions. Finally, using a mark-recapture experiment, we discovered a strong temporal segregation in patrolling activity of males from two co-mimetic sister species. Such divergence in phenology may favour sympatry between closely-related species, despite behavioural interferences induced by the local convergence in colour pattern. Altogether, our findings show that temporal segregation may facilitate the co-existence of closely-related species sharing the same ecological niche, suggesting that phenological shifts may represent an overlooked factor of sympatric speciation. Our study therefore highlights how the evolution of multiple traits may favour species diversification in sympatry by partitioning niche in different dimensions.

## Introduction

Natural communities are composed of multiple species involved in diverse ecological interactions either facilitating or impairing their co-existence. Shared ancestry induces ecological and morphological similarity of closely-related species, promoting their co-existence in similar niches (Wiens and Graham, 2005). However, such similarity also inherently limits this co-occurrence (Abrams, 1983; MacArthur and Levins, 1967) by increasing interspecific competition and/or reproductive interference (Brown and Wilson, 1956; Grant and Grant, 2006; Gröning and Hochkirch, 2008; Kyogoku, 2015). Furthermore, gene flow is more likely to happen between closely-related species during secondary contacts (Hewitt, 2000), and may impair their genomic and phenotypic divergence (Bolnick and Fitzpatrick, 2007). The genomic and ecological similarity may thus limit co-existence of closely-related species in sympatry.

Some types of genetic exchanges and ecological interactions might nevertheless facilitate such co-existence. Gene flow between closely-related species can indeed enhance local adaptation through introgression, enabling the rapid emergence of locally adaptive traits in colonizing species (Dasmahapatra et al., 2012; Huerta-Sánchez et al., 2014; Vekemans, 2010). Commonly inherited traits may locally benefit to individuals of closely-related species: when facing common predators for example, responding to alarm cues emitted by heterospecifics (Chivers et al., 2002; Dalesman and Rundle, 2010), or sharing a common warning signal with co-occurring species (Müller, 1879; Sherratt and Beatty, 2003). Such interspecific positive dependence is predicted to favour species co-existence (Aubier, 2020).

Shared signals among species may however enhance interspecific behavioural interference (Grether et al., 2017), including heterospecific courtship and male-male competition (Gröning and Hochkirch, 2008). Close relatedness between species then often results in reinforcement mechanisms promoting either divergent evolution of other mating cues (Saetre et al., 1997; Smadja and Ganem, 2008) or cryptic signalling (Dasmahapatra et al., 2012; Fordyce et al., 2002). Alternatively, shared signal among species may induce interspecific territoriality (Drury et al., 2020; Grether et al., 2017; Souriau et al., 2018), thereby reducing resource competition and facilitating coexistence in segregating spatial niches within the geographic areas (e.g. Souriau et al., 2018; Tobias and Seddon, 2009). This spatial segregation might be especially promoted in recently diverged species where heterospecific competition and mating are high (Drury et al., 2020; Drury et al., 2015). Phylogenetic relatedness and phenotypic resemblance are indeed strong predictors of interspecific territoriality (Losin et al., 2016). Ecological factors preventing character divergence might therefore also promote interspecific territoriality (Drury et al., 2020).

Selection promoting traits convergence is particularly strong in mimetic butterflies, therefore limiting population divergence in sympatry. Convergence in warning pattern indeed frequently involves convergence in other traits like flight height (Elias et al., 2008) or host-plant (Willmott and Mallet, 2004), because sharing a common microhabitat induces a higher similarity of predator communities encountered and therefore an enhanced protection (Gompert et al., 2011). Nevertheless, the benefits conferred by overlapping visual signals and ecological niches in mimetic species may in turn incur fitness cost through increased heterospecific rivalry, courtship and the expression of Dobzhansky-Müller incompatibilities in hybrids (Estrada and Jiggins, 2008; Mérot et al., 2015; Welch, 2004), ultimately limiting the co-existence of closely-related mimetic species in sympatry (Aubier et al., 2017). The conflicting ecological interactions between mimetic species therefore question their persistence in sympatry when they are closely-related.

Here we investigate how territoriality and cryptic signalling might favour the coexistence of closely-related mimetic species by focusing on sympatric *Morpho* butterfly species, displaying striking large blue iridescent wings (Debat et al., 2018). Although not chemically defended, these butterflies are very difficult to capture because of their fast, erratic flight. The contrast between their dorsal bright blue and ventral cryptic brownish wing surfaces induces a flash pattern during flapping flight, that was suggested to confuse predators, further increasing their difficulty of capture (Debat et al., 2018; Murali, 2018; Pinheiro and Campos, 2019; Young, 1971). Predators may then learn to avoid such elusive prey harbouring the conspicuous blue patterns. Some sympatric *Morpho* species were shown to locally converge in their blue pattern, probably because of frequency-dependent selection generated by predator behaviour, in a similar way as Müllerian mimics (Llaurens *et al.* 2020). The iridescent blue colouration of *Morpho* butterflies shared by sympatric species may thus reduce individual predation by advertising escape ability (the “escape mimicry” hypothesis, see Páez et al., 2020). Such local convergence might in turn impair species recognition and mate choice. This duality questions how closely-related co-mimetic species become sexually isolated and may coexist in sympatry.

## Results and Discussion

We focused on a single locality where three mimetic species from the genus *Morpho (M. achilles, M. helenor* and *M. deidamia)* live in sympatry. In this field site, males from these three closely-related species display their typical patrolling behaviour along the river bed, allowing investigating species interactions in sympatry, as well as genomic exchanges among sympatric species.

### Closely-related *Morpho* species with mimetic colouration live in sympatry

We performed a series of experiments at this field site, located in the Amazonian Peru, at the foothill of the Andes. We first estimate the local abundances of the different *Morpho* species at the study site using a mark-recapture experiment (see material and methods and Fig. S1). *Morpho achilles* was the most abundant species (mean ± se = 264 ± 68), followed by *M. helenor* (mean ± se = 195 ± 49). Individuals from *M. deidamia* were too rarely caught to allow estimating population size, suggesting a lower abundance at this site (Table S1). These three mimetic species thus co-occur at the study site, allowing relevant experiments on species co-existence.

### Heterospecific interactions lead to reproductive interference among sympatric species

To test whether similarity in colour pattern among these sympatric *Morpho* species leads to heterospecific rivalry and courtship, we then investigated the response of patrolling males to butterfly dummies placed in the field. We built realistic dummies using the actual wings of captured *Morpho*, set up on a solar-powered fluttering device reproducing butterfly flying behaviour (Fig. 1; Movie S1; see also Fig. S2, S3). To ensure that only visual cues were triggering the interactions, the wings were washed in hexane prior mounting, ruling out any effect of pheromones (Darragh et al., 2017). We built 10 different dummies with the wings of specimens from different species and sexes, all caught in the same Peruvian site. We used both sexes of the two mimetic sister-species *M. helenor* and *M. achilles*, as well as of a third mimetic species *M. deidamia*, that all present an iridescent blue band bordered by proximal and distal black areas, and the phenotypically distinct *M. menelaus,* exhibiting fully blue iridescent wings (local dummies: *n* = 8) (Fig. 2, S2). The last two dummies were built from the wings of males *M. helenor* and *M. achilles* captured in French Guiana (exotic dummies: *n* = 2), exhibiting a narrower blue band relative to local – Peruvian – individuals (Fig. 2, S2). We tested all dummies (n=10) following a randomized design: a different dummy was placed each morning at the same site and left fluttering on the river bank for 5 hours. This was replicated 4 times per dummy.

**Fig. 1.**
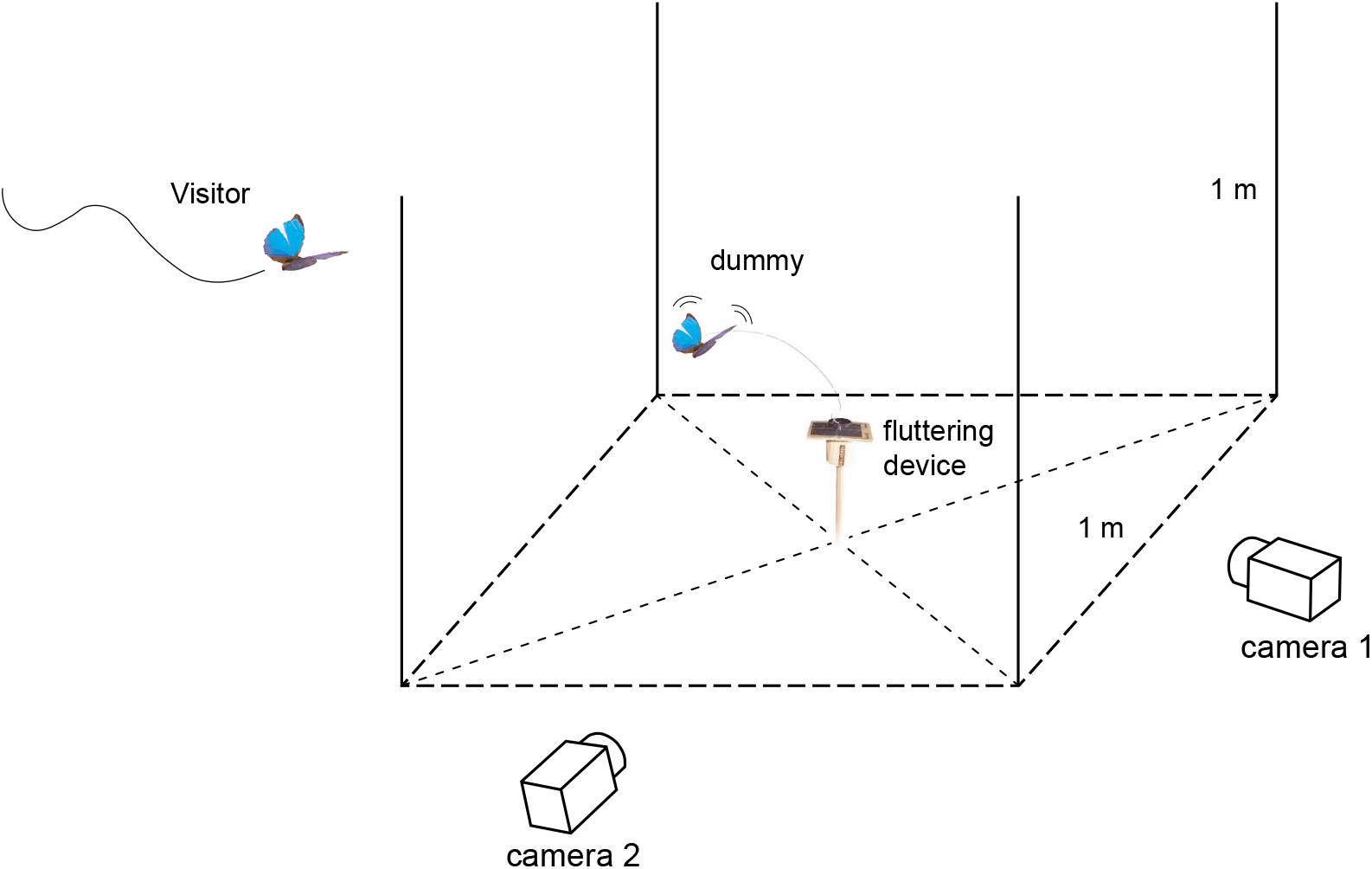
Experimental set up used to study flight interaction. The dummy butterfly was placed at the centre of a cubic area materialized by four 1 m3-sticks, and fixed to a solar-powered fluttering device reproducing butterfly flying behaviour. Interaction between visitor and dummy butterfly (defined as a visitor entering in the cubic area) were recorded using stereoscopic high-speed videography system, allowing to quantify flight trajectory during the interaction (see Fig. 3).

**Fig. 2.**
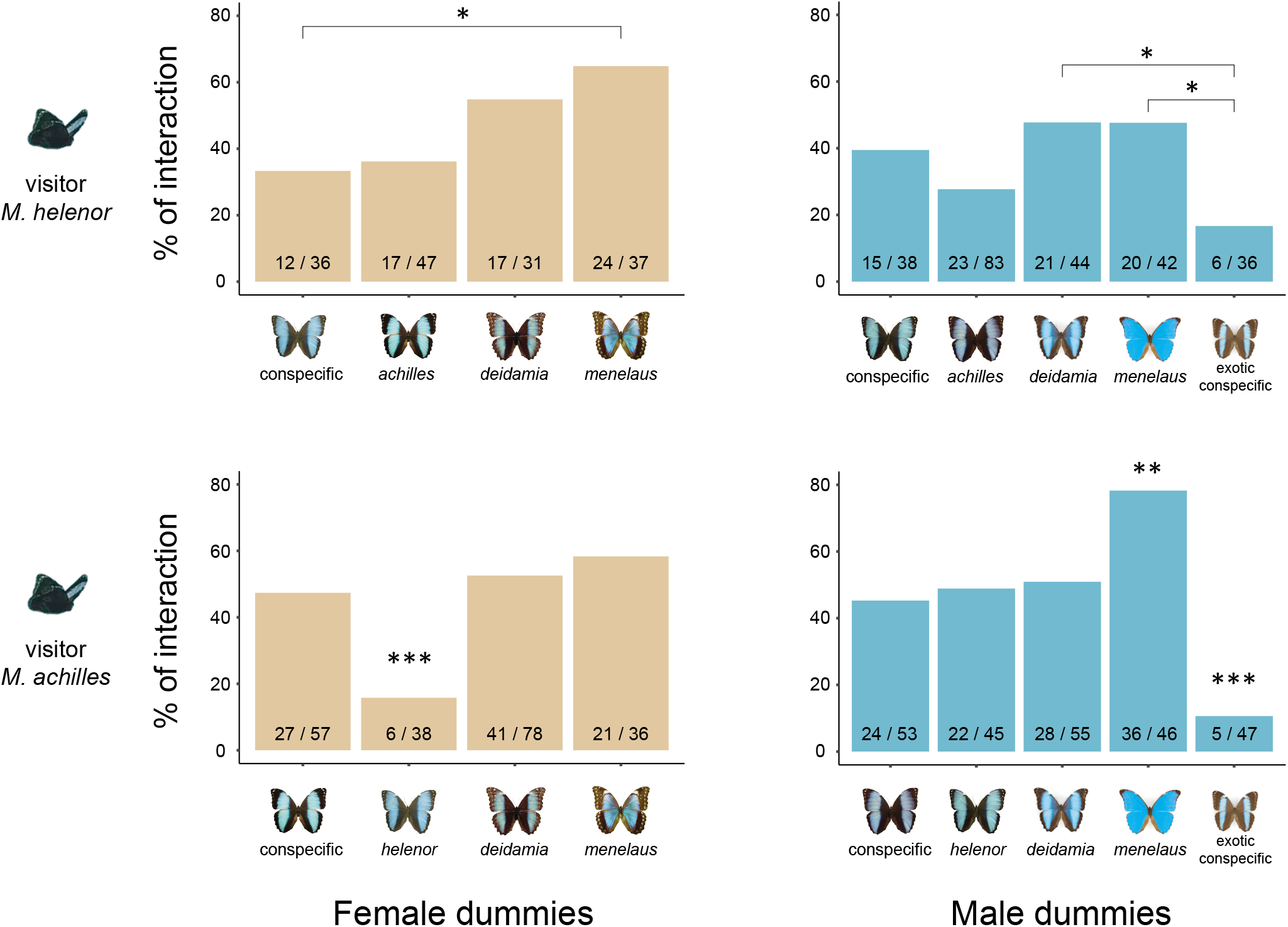
Variation in interaction frequency with conspecific and congener dummies in two mimetic sister *Morpho* species. *Morpho helenor* (top row) and *M. achilles* (bottom row) engage in interaction with sympatric conspecifics and congeners (females: left column, males: right column) in broadly similar proportion. Note that males *M. achilles* showed limited courtships with the female of its co-mimetic sister and that percentage of interaction was consistently lower with the exotic male conspecific in both species. Raw data « nb of interactions / nb of approaches » are indicated on each barplot.

Over a two months period, we recorded 2,700 patrols of wild males and studied behavioural responses to dummies. We specifically focused on the behaviour of butterflies from the two mimetic sister species *M. helenor* and *M. achilles,* which represented the large majority of the passing males (35% and 47% respectively Fig. S4). During these sessions, we scored two behaviours: (1) *approaches,* when a marked change in the trajectory of the passing butterfly towards the dummy was observed and (2) *interactions*, when the butterfly entered a 1m^3^ zone around the dummy (Fig. 1). This procedure allowed us to test whether patrolling *Morpho* males (1) are more strongly attracted by the colour pattern of their conspecifics as compared to that of other species, (2) discriminate between sexes and (3) are more attracted by local colour patterns than by exotic ones.

#### Long-range visual attraction to blue butterflies

About half of the patrolling individuals deviated from their flight path to *approach* the setup (see Fig. S5, S6 and Table S2,S3 for more details). The differences in wing area and proportion of iridescent blue on the wing among the dummies had a limited effect on the percentage of *approach* (Fig. S7 and Table S4). The long range blue signals emitted by the different wing patterns displayed in the different species tested thus appeared similarly attractive for patrolling males of the sympatric species *M. helenor* and *M. achilles*, suggesting they may approach to anything roughly recognized as a potential mate or rival.

#### Significant discrimination between local and exotic butterflies

In both *M. helenor* and *M. achilles*, about 40% of the *approaches* resulted in an *interaction*, the visitor typically flying in circles around the dummy, with little evidence of discrimination between sympatric conspecifics and congeners, as well as between female and male dummies (Fig. 2). No effect of sex or identity of the dummy was detected on the duration of the *interactions* (Kruskal-Wallis test, dummy sex: Chi-square = 3.71, *P* = 0.20, df =1; dummy identity: Chi-square = 14.03, *P* = 0.12, df =9). However, *interactions* occurred in markedly lower proportions with the exotic dummies as compared to most local dummies (Fig. 2). Males *Morpho* do discriminate between slightly different colour patterns (*i.e.* large blue band in local individuals, vs. narrower blue band in exotic ones), but yet they largely engage in *interactions* with congeners bearing locally known signals, including signals sharply dissimilar from their own (*e.g.* fully blue wings in *M. menelaus*). Overall, dummies with larger wing area and greater proportion of iridescent blue colouration were more likely to trigger *interaction* (Fig. S7), suggesting that these characteristics are used by patrolling males to discriminate among dummies. Such a stronger response to local, known phenotypes (including heterospecifics) than to exotic ones (including conspecifics), has been suggested to reflect high interspecific territoriality, particularly in males (e.g. Tobias and Seddon, 2009). Consistent with this hypothesis, *interactions* were the most indiscriminate among male local dummies, as compared to female dummies (Fig. 2).

#### Recognition of conspecific females in Morpho achilles

While no significant differences were observed in *interactions* with dummies displaying male wings of *M. achilles* and *M. helenor*, patrolling *M. achilles* males clearly avoided its co-mimetic *M. helenor* female, but interacted indiscriminately with the dummies of the other sympatric species (i.e. *M. deidamia* and *M. menelaus*) (Fig. 2). Such a specific, acute visual discrimination towards its phenotypically closest sister species suggest that mate-recognition based on cryptic signals could have evolved in *M. achilles*, possibly as a result of reinforcement selecting against hybridization (Servedio and Noor, 2003). Reinforcement process may also occur through divergence in olfactory cues enabling discrimination among species (Mérot et al., 2015; Smadja and Ganem, 2008), although this remains to be investigated in *Morpho*.

To test whether the behaviour of *M. achilles* males differs when interacting with conspecific male and female dummies, we then equipped our set-up with a stereoscopic high-speed videography system, enabling to quantify the three-dimensional flight kinematics of visiting butterflies in natural conditions on a sub-set of sessions (Fig. 1). Striking behavioural differences were observed between the interactions toward males and females (*n* = 14 flights analysed for each sex): on average, wild males circled closer to the female than to the male dummy (regression of proportion of time spent *vs*. distance from dummy female: *P* < 0.001; *R*^2^ = 0.07, dummy male: *P* = 0.09; *R* = 0.01; Fig. 3). Besides, males approached the female dummy following a smoothly decelerating flight path, ensuring a steady speed when close to the female, whereas they showed more erratic accelerations around the male dummy (regression of acceleration on distance from dummy female: *P* < 0.001; *R^2^* = 0.14, dummy male: *P* = 0.07; *R^2^* = 0.02; Fig. 3). During these aerial *interactions*, *M. achilles* males adjust their flight behaviour according to the sex of their conspecific, relying solely on colour pattern visual cues.

**Figure 3.**
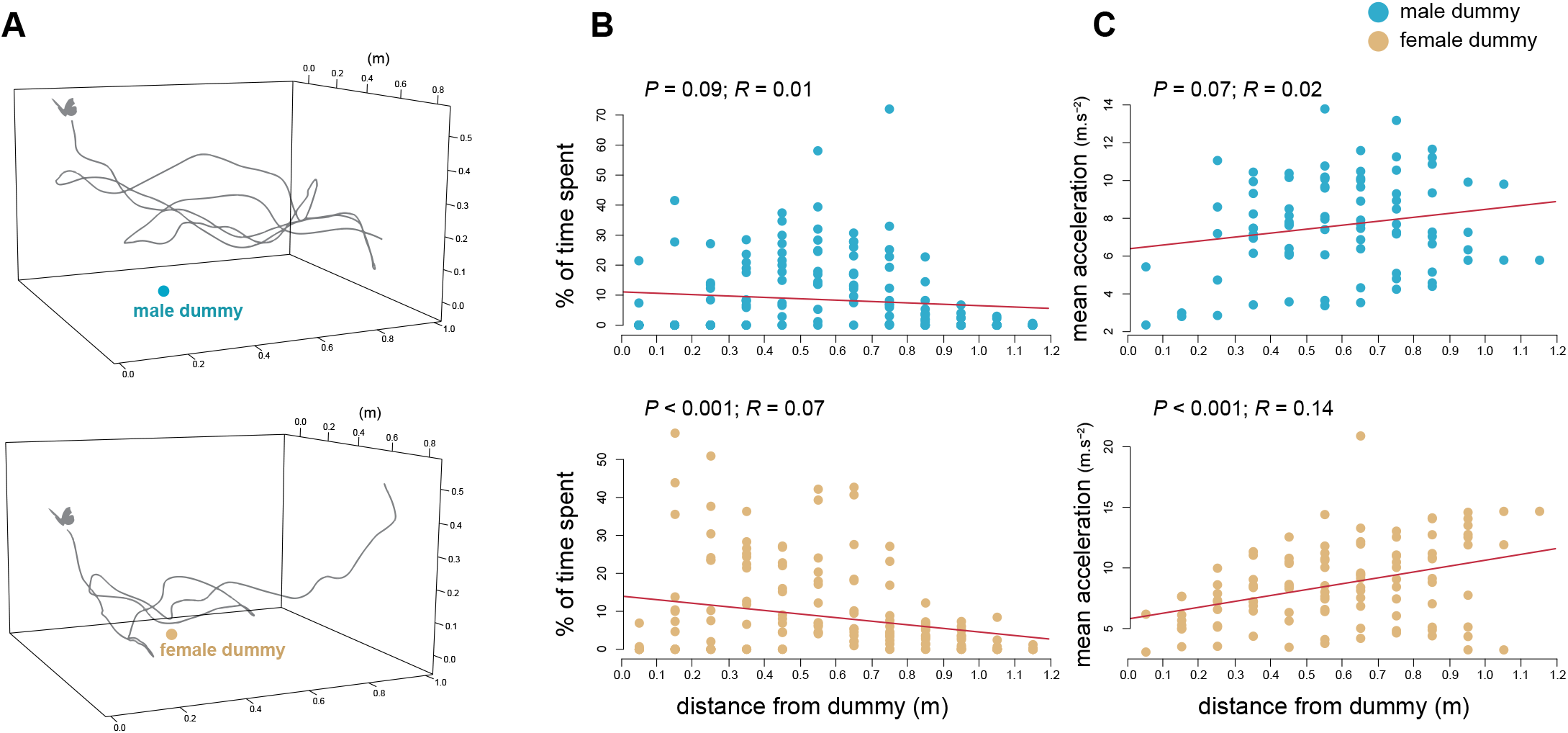
Three-dimensional kinematic of contest and courtship flights. (**A**) Example of flight path when circling around the dummy conspecific male (top) and female (bottom). (**B**) The proportion of time spent circling at short-distance from the dummy significantly increases for dummy female only. (**C**) A significantly positive relationship between acceleration and distance from dummy is observed for dummy female only. N = 14 flight interactions analysed with the dummy male and 14 with the dummy female.

#### High reproductive interferences and marked differences in discrimination behaviour between closely-related species

Altogether, our results thus clearly suggest that heterospecific contests frequently occur among males from sympatric *Morpho* species, and that heterospecific mating or mating attempts can frequently occur, leading to strong reproductive interferences in those sympatric species. Males from the species *M. helenor* were found to approach dummies significantly more than *M. achilles* (mean % of approach in *M. helenor* = 48.3 ± 15.7; *M. achilles* = 39.7 ± 14.8; *P* = 0.01). They did not showed increased *interactions* with conspecific females as compared to *M. achilles* females (Fig. 2). Behavioural interferences between these sympatric species are thus large, with a higher indiscriminate behaviour in *M. helenor* than in *M. achilles*. These interspecific interactions would then induce high costs to both sexes: indiscriminate aerial contest performed with all passing individuals are energetically costly to males (Kemp, 2013; Takeuchi, 2017), while females are likely harassed by such indiscriminate mating attempts (Gröning and Hochkirch, 2008; Kyogoku, 2015; Kyogoku and Nishida, 2013). These costs could be higher in *M. helenor* where discrimination capacities were found lower than in *M. achilles*. Indiscriminate mate-searching behaviour in *M. helenor* moreover likely increases competition for females among males from sympatric species. This is expected to promote interspecific territoriality (Drury et al., 2020; Grether et al., 2017), in agreement with the frequent heterospecific contests observed between males *M. helenor* and *M. achilles*. These costly behaviours question the stable coexistence and the reproductive isolation of sympatric *Morpho* species.

### Limited genetic exchanges between species despite close relatedness and reproductive interference

We then sampled DNA from 31 butterflies caught at this site (13 *M. achilles*, 8 *M. deidamia* and 10 *M. helenor*), to explore the level of genomic exchanges among species. To specifically test whether the genomic patterns of polymorphism (Fig. S8, S9) and divergence (Fig. S10) reflect episodes of introgression between species, we performed RAD-sequencing and statistically evaluate alternative scenarios of speciation (Fig. S11), with and without gene flow using the demographic inferences with linked selection (DILS) approach (Fraisse et al., 2020). By comparing in a hierarchical approach eight categories of models according to their temporal patterns of migration between *M. helenor*, *M. achilles* and *M. deidamia*, this Approximate Bayesian Computation approach provides strong statistical support for an isolation of *M. deidamia* with both *M. helenor* and *M. achilles* (Fig. 4; posterior probability = 0.91). The best scenario among those proposed also describes a divergence between *M. helenor* and *M. achilles* with migration restricted to the first generations after the split (posterior probability = 0.82). This analysis suggests that current putative hybridizations would not represent a significant source of intraspecific genetic diversity (Fig. 4; Table S5). Consistent with the phylogeny (Chazot et al., 2016), demographic inferences performed on our Peruvian populations revealed more recent time of split between the sister species *M. achilles* and *M. helenor* (*T_2_* = 1.11 million generations) than between these two species and *M. deidamia* (*T_1_* = 4.13 million generations). This strict isolation despite close relatedness and reproductive interference between *M. achilles* and *M. helenor* questions the ecological factors limiting hybridization in sympatry.

**Fig. 4.**
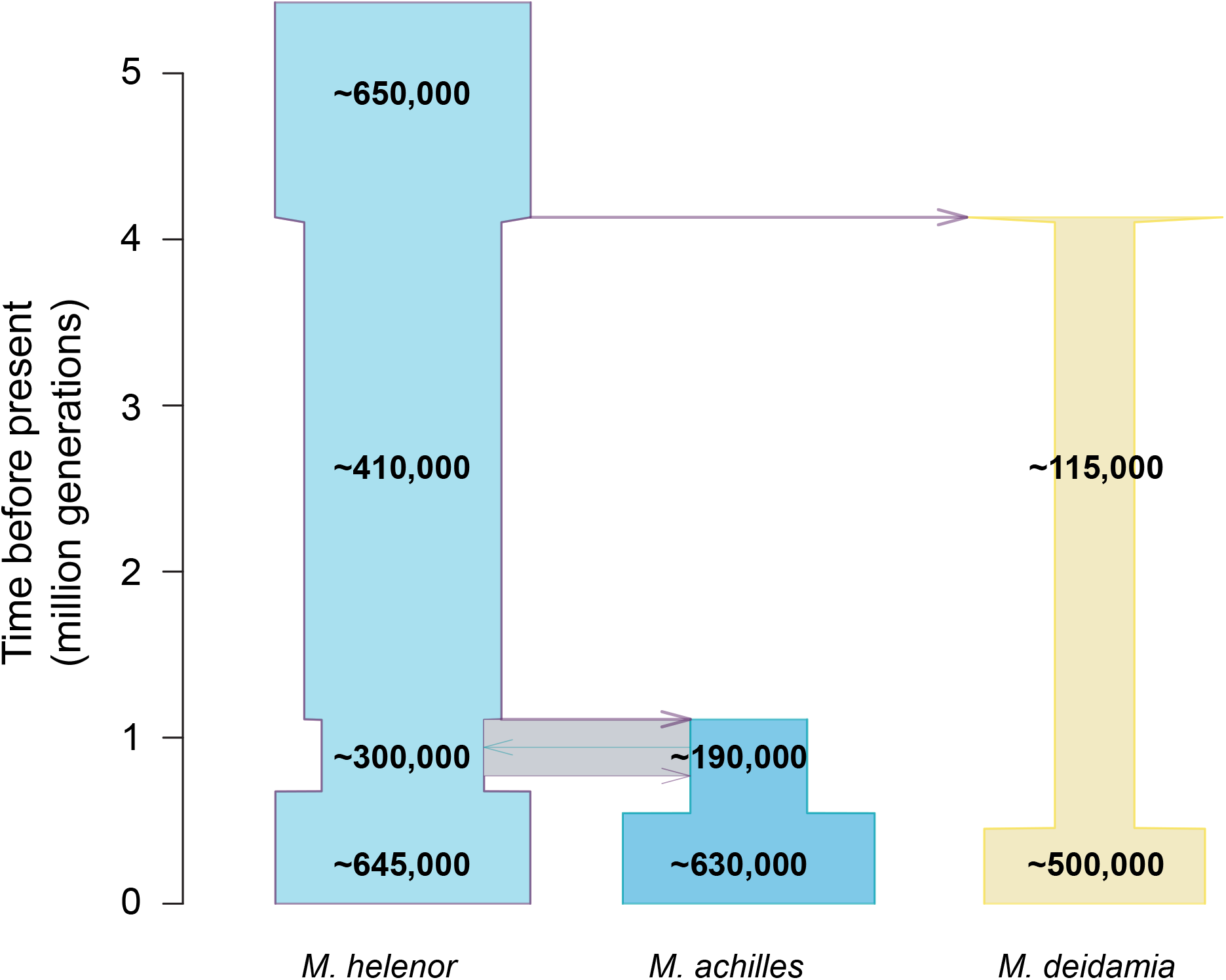
Best inferred demographic scenario from RAD-sequencing data. Eight categories of scenarios were compared according to different temporal patterns of gene flow: between *M. helenor* and *M. achilles* (ancestral migration or secondary contact); with *M. deidamia* (strict isolation, migration only with *M. helenor*, migration only with *M. achilles*, migration between the 3 species). The times of demographic events (speciation, cessation of migration and changes n population size) are shown on the Y-axis in millions of generations. The grey rectangle indicates the period when *M. helenor* and *M. achilles* were, according to the demographic model fitting the best the molecular dataset, genetically connected through migration events.

### Temporal segregation between sympatric sister-species

By analysing the temporal variations in the mark-recapture experiments, we observed a striking difference in patrolling time among species (Kruskal-Wallis test: Chi-square = 179.7, *P* < 0.001, df = 3), with little overlap between the sister species *M. achilles* and *M. helenor* (Fig. 5). Males *M. helenor* patrolled earlier than *M. achilles* (mean patrolling time ± s.d. = 11:14 ± 00:45 vs. 12:35 ± 00:40, respectively). Patrolling time in *M. deidamia* (12:40 ± 00:46) however fully overlaps that of *M. achilles*. Time of capture was remarkably similar among recaptures of a same individual in the species *M. achilles* (correlation between time of first *vs*. second capture: *r* =0.40; *P* = 0.05), suggesting a regularity in patrolling time at the individual level in this species (Fig. 5). Whether such individual temporal regularity is genetically determined or reflects a plastic behaviour (Groot, 2014; Schöfl et al., 2009) remains to be investigated. The close phenotypic similarity of sympatric *Morpho* species, probably promoted by escape mimicry (Llaurens *et al.* 2020) might thus enhance reproductive interferences between then closely-related species *M. helenor* and *M. achilles* and may have favoured the evolution of divergent temporal niches. Reproductive and/or aggressive interference are generally expected to promote spatial or temporal habitat segregation between species because it reduces the cost of negative interspecific interactions (Grether et al., 2017; Robinson and Terborgh, 1995).

**Fig. 5.**
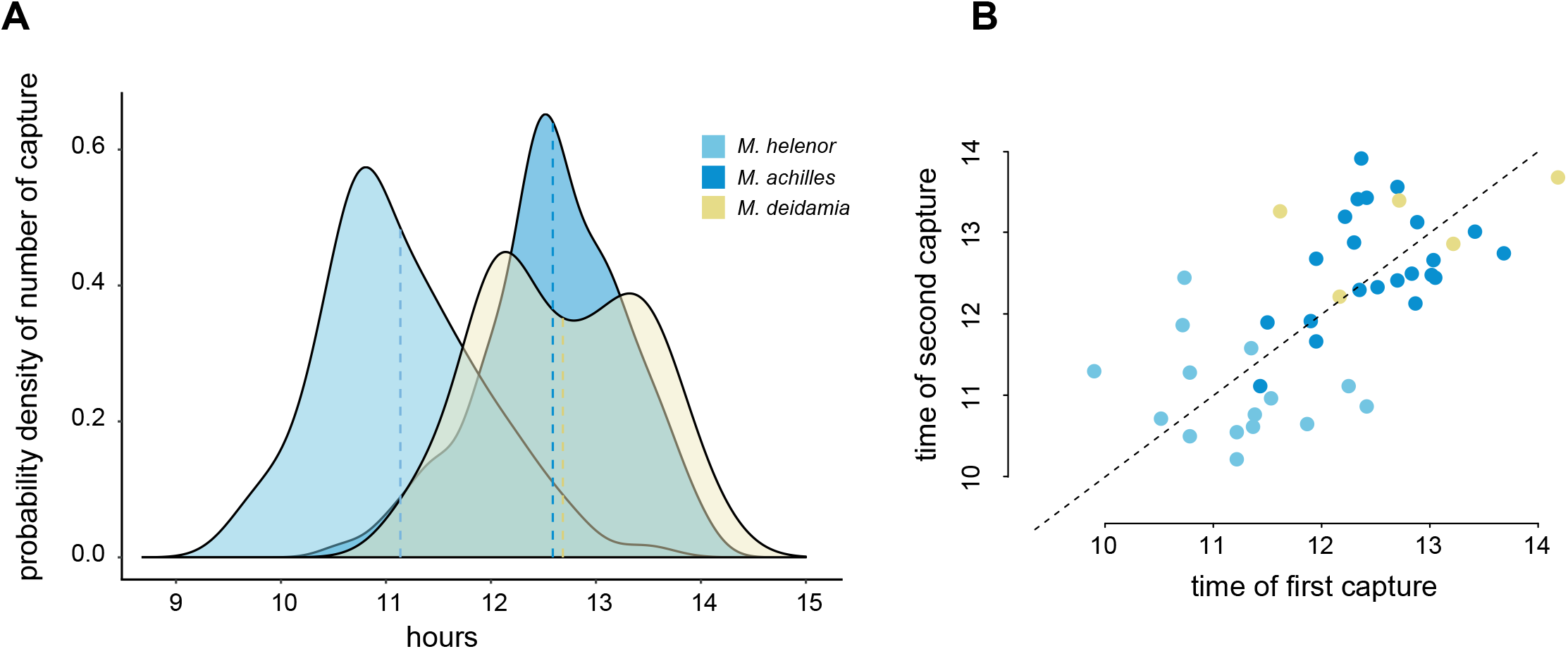
Patrolling time among sympatric species and individuals. (A) Segregation of patrolling time over the day among *Morpho* species. Dashed vertical lines indicate the mean flight time. (B) Plot of first versus second capture time among recaptured individuals. Dashed line represents exact same time between first and second capture.

Altogether, our results suggest that a strong interspecific competition occurs among males from mimetic *Morpho* species in sympatry, but that this competition is likely mitigated by their temporal segregation. Heterospecific courtships may also be limited, if the temporal segregation observed in *Morpho* males is mirrored by a similar temporal partitioning of females activities. The activity of females butterflies is often challenging to estimate because of their highly cryptic behaviour (Devries et al., 2008), and was not measured in our study, as they were too rarely encountered. Synchronization of mating activity between sexes is however likely (Hirota et al., 2001; Iwasa and Obara, 1989), because any deviation of males relative to females activity would reduce the probability of intra-specific mating (Groot, 2014; Schöfl et al., 2009). In contrast, a shift in mating timing among species can act as a powerful isolation mechanism (Taylor and Friesen, 2017), and might explain how co-mimetic *Morpho* species can coexist in a same habitat while remaining sexually isolated.

Temporal segregation in butterfly mating activities may be a widespread process enabling the persistence of diversity-rich assemblages, as suggested by several reports of temporally structured sexual activities in other butterflies (Callaghan, 1982; Freitas et al., 1997; Kemp and Rutowski, 2001), including closely-related species (Devries et al., 2008). Our study on mimetic butterflies highlights that the co-existence of closely-related species can generate complex ecological interactions, both mutualistic (mimicry) and antagonistic (reproductive interference), that could be mitigated by shifts in temporal niches. Our study therefore highlights how the evolution of multiple traits may favour species diversification in sympatry by partitioning niche in different dimensions.

## Material and Methods

### Study site and population

The study was conducted between July and October 2019 in the North of Peru. We focused on populations of coexisting *Morpho* species present in the regional park of the Cordillera Escalera (San Martin Department) near the city of Tarapoto. Both the capture-recapture and the dummy experiment were performed at the exact same location, on the bank of the Shilcayo river (06°27′14.364″ S, 76°20′45.852″ W).

### DNA extraction and RAD-Sequencing

We used 31 wild males caught on this sites to perform population genomics analyses (*M. achilles* - *n* = 13, *M. helenor* - *n* = 10 and *M. deidamia*-*n* = 8). DNA was then extracted from each sample using a slice of the thorax, using Qiagen kit DNeasy Blood & Tissue. DNA quantification (using Qubit and Nanodrop methods) and DNA quality (by electrophoresis) was performed before the sequencing carried out in the MGX-Montpellier GenomiX platform (Montpellier, France). DNA digestion was performed using the Pst1 enzyme and paired-end RAD-sequencing has then been performed in one run as set out by Baird and Etter (2008) (2008), giving 299 billion reads, comprising R1 and R2 reads for each sequenced fragment. The sequencing has been performed on Illumina sequencers HiSeq2500 so that reads (125bp) were expected to be of high quality, without missing base (N content). MGX then demultiplexed the data using the software Stacks (Catchen et al., 2013; Catchen et al., 2011), allowing assigning each read to his sample ID. Adapters have all been correctly removed from each reads.

### Reads quality, alignment and data set generation

Reads quality has been performed using FastQC v0.11.9 software (http://www.bioinformatics.babraham.ac.uk/projects/fastqc/). All reads have the same length: 119bp and 125bp for R1 and R2 paired-end reads respectively. Per base sequence quality were high for R1 paired-end sequences (>36) and for R2 paired-end sequences (>32) with a quality score per sequence around 39 for each reads (40 being the maximum). Sequence content per base was accurate for both R1 and R2 paired-end reads. GC content per was slightly higher than the theoretical distribution (calculated with the mean GC rate) for each reads. Thus, the high quality of reads allows avoiding read trimming or deletion.

One read alignment was realized using the Stacks V2.5 software (http://catchenlab.life.illinois.edu/stacks/). Parameters have been set using r80 methods which maximize the number of SNPs or loci shared by at least 80% of the samples (Paris et al., 2017). The optimized parameters are “max distance between stacks” (inside each sample) and “number of mismatches between stacks” (between samples). Every other parameters have been kept to default values. Data sets were in the form of VCF data file (containing all the SNPs found in the alignment) and fasta data file (which contain the two alleles found for every loci for each sample). A RAD-sequencing locus is thus a sequence composed of 1 to 3 paired-end reads aligned together and delimited by restriction sites.

To run DILS-ABC inferences, Stacks fasta file has been converted in another fasta file compatible with DILS (https://github.com/CoBiG2/RAD_Tools).

### Demographic inferences

Eight categories of demographic models were compared, according to temporal patterns of introgression. This was done to answer two questions on gene flow in *Morpho*: 1) is there ongoing migration between *M. helenor* and *M. achilles*? 2) do *M. helenor* and/or *M. achilles* exchange alleles with *M. deidamia*? This was achieved by an ABC approach using a version of *DILS* adapted to samples of three populations/species (Fraisse et al., 2020).

A generalist model was studied (Fig. S11). This model describes an ancestral population subdivided in two populations: the ancestor of *M. deidamia* and the common ancestor of *M. helenor/M. achilles*. The latter population was further subdivided into the three species/populations currently sampled. Each split event is accompanied by a change in demographic size, the value of which is independent of the ancestral size. In addition, given clear genomic signatures for recent demographic changes with largely negative Tajima’s *D*, we implemented variations for the effective sizes of the 3 modern lineages at independent times. Finally, migration can occur between each pair of species/populations. Migration affecting the *M. helenor/M. achilles* pair can either be the result of secondary contact after a period of isolation (ongoing migration), or of ancestral migration (current isolation) as in (Roux et al., 2016).

As this model is over-parameterized, our general strategy is to investigate the above two questions by comparing variations of this generalist model. Thus, to test the gene flow between *M. helenor* and *M. achilles*, we compare two categories of models. 1) With random parameter values for all model parameters including the ongoing migration between *M. helenor* and *M. achilles* (gene flow resulting from a secondary contact between them); 2) as above, but with the migration between *M. helenor* and *M. achilles* set to zero after a randomly drawn number of generations following their split.

For each models, 50,000 of simulations using random combinations of parameters were performed. Parameters were drawn from uniform prior distributions. Population sizes were sampled from the uniform prior [0-1,000,000] (in diploid individuals); the older time of split was sampled from the uniform prior [0-8,000,000] (generations); ages of the subsequent demographic events were sampled in a uniform prior between 0 and the sampled time of split. Migration rates 4.*N.m* were sampled from the uniform prior [0-50]. Both migration rates and effective population sizes are allowed to vary throughout genomes as a result of link selection following (Charlesworth et al., 1993; Cruickshank and Hahn, 2014; Roux et al., 2014).

On each simulated data set we calculate a vector of means and standard deviations for different summary statistics: intraspecific statistics (π for *M. helenor*, π for *M. achilles*, π for *M. deidamia*, θw for *M. helenor*, θw for *M. achilles*, θw for *M. deidamia*, Tajima’s *D* for M. helenor, Tajima’s *D* for *M. achilles*, Tajima’s *D* for *M. deidamia*) and interspecific statistics (gross divergence, net divergence and FST for all three possible pairs; ABBA-BABA *D*).

Statistical comparisons between simulated and observed statistics were performed using the R package *abcrf* (Pudlo et al., 2014; Raynal et al., 2019).

### Mark-recapture experiment

To estimate abundance and patrolling activity among *Morpho* species, we performed capture-mark-recapture between 9 a.m. and 2 p.m. (flight activity in *Morpho* is drastically reduced in the afternoons at this site) during 17 sunny days. Although on few days, capture was aborted because of bad weather annihilating butterfly activity, the 17 capture sessions were mostly consecutives, as they were performed in a 22 days period. All butterflies were captured with hand-nets, identified at the species level, and numbered on their dorsal wing surface using a black marker. The exact time of each capture was annotated. Butterflies captured while inactive, such as those laying on a branch or on the ground were excluded from the analysis to focus exclusively on actively patrolling individuals. We measured patrolling time for a total of 295 occasions, including 78 recaptures (i.e. 217 individuals were captured at least once). All captured individuals were males. Individuals *M. achilles* were the most frequently captured (*n* = 121), followed by *M. helenor* (*n* = 95). Individuals *M. deidamia* were about half less captured (*n* = 48), and individual *M. menelaus* were the least captured (*n* = 34). Based on capture-recapture histories, we estimated individual abundance for each species using a loglinear model implemented in the R package *Rcapture* (Baillargeon and Rivest, 2007) (Fig. S1). Given the short duration the sampling period (22 days) relative to the longevity of adult *Morpho* butterflies (several months, Garcia et al., 2014), we used a closed-population model assuming no effect of births, deaths, immigration and emigration. Abundance was estimated in *Morpho helenor* and *M. achilles* only, as capture and re-capture events were too few in the other species (*M. deidamia* and *M. menelaus*) to allow estimating population size (Table. S1). Because striking differences in patrolling time were rapidly observed among *Morpho* species, we used time of the day as a predictor of species identity in order to distinguish between *M. helenor* and *M. achilles* in the below-described experiment because butterflies from these two species are morphologically too similar to be identified while flying (Fig. S12). After the 17 nearly-consecutive days of capture, one day of capture was repeated every 2 weeks during 2 months in parallel to the dummy experiment (described below), to verify that temporal activity was stable over time (Fig. S12).

### Dummy butterflies experiment

We investigated the response of patrolling males to sympatric conspecifics, congeners and of exotic conspecifics, using dummies placed on their flight path. Dummies were built with real wings dissected and washed with hexane to remove volatile compounds and cuticular hydrocarbons, ensuring to test only the visual aspect of the dummies. We mounted the wings on a solar-powered fluttering device (Butterfly Solar Héliobil R029br) that mimics a flying butterfly, thereby increasing the attractiveness of the dummy. The fluttering dummy was positioned on the river bank, and placed at the centre of a 1m^3^ space delimitated with four vertical stacks (Fig. 1). Patrolling *Morpho* butterflies that deviated from their flight path to approach the dummy but did not enter the cubic space were categorized as *approaching*. Any *Morpho* butterfly entering the cubic space was considered as *interacting* with the dummy. Those passing without showing interest to the setup were categorized as *passing*. All patrolling individuals were identified at the species level, either visually for *M. menelaus* and *M. deidamia*, or based on time of the day for *M. helenor* and *M. achilles* (Fig. S12). By continuously filming the setup using a camera (Gopro Hero5 Black set at 120 images per second) mounted on a tripod, we also measured the duration of the interactions (*i.e.* the time spent in the cubic space) occurring between patrolling male and the dummy. The ten dummies were each tested during 4 sunny days from 9 a.m. to 2 p.m. (*i.e.* during 5 hours). This resulted in 40 days of experiment over which each dummy was left fluttering on the river bank for a combined duration of 20 hours. Dummies were randomly attributed to each day of experiment.

### Three-dimensional kinematics of flight interaction with the dummies

To test if *Morpho* males shows different flight behaviours when interacting with male and female dummy, we filmed the flight interactions using two orthogonally positioned video cameras (Gopro Hero5 Black, recording at 120 images per second) around the dummy setup (Fig. 1). Stereoscopic video sequences obtained from the two cameras were synchronized with respect to a reference frame (here using a clapperboard). Prior to each filming session, the camera system was calibrated with the direct linear transformation (DLT) technique (Hartley and Zisserman, 2003) by digitizing the positions of a wand moved around the dummy. Wand tracking was done using DLTdv8 (Hedrick, 2008), and computation of the DLT coefficients was performed using easyWand (Theriault et al., 2014). After spatial and temporal calibration, we also used DLTdv8 to digitize the three-dimensional positions of both the visiting (real) butterfly and the dummy butterfly at each video frame by manually tracking the body centroid in each camera view. Butterfly positions throughout the flight trajectory were post-processed using a linear Kalman filter (Muijres et al., 2014), providing smoothed temporal dynamics of spatial position, velocity and acceleration of the body centroid. Based on these data, we investigated how spatial position, speed and acceleration of the visitor butterfly varied over the course of the interaction. We proceeded by dividing space into 10 cm spherical intervals around the dummy position ranging from 0 to 1.2 meters distance, and computed the proportion of time spent, the mean speed and acceleration of the interacting butterfly within each distance interval (Fig. 3). We analysed a total of 28 interactions performed by individual *Morpho achilles* male, including 14 with the dummy of its conspecific male and 14 with the dummy of its conspecific female. Analysed interactions lasted in average 1.44±0.87 (mean±sd) seconds.

### Statistical analysis of behavioural experiments

Differences in patrolling time were assessed by testing the effect of species on time of capture using Kruskal-Wallis test. To test the effect of visitor identity and dummy characteristics on the number of approaches and interactions, we performed logistic regressions. *Approach* was treated as a binary variable, where 0 meant “passing without approaching” and 1 meant “approaching the dummy setup”. For the interactions, we only considered individuals approaching the setup, such as 0 meant “approaching without entering the cubic space” and 1 meant “entering the cubic space”. This allowed getting rid of the uncertainties on whether passing individuals had actually seen the setup or not. We first tested the effect of visiting species on *approach* and *interaction* while controlling for dummy’s characteristics to test for intrinsic differences in territoriality (or ‘curiosity’) among species. We then tested the effect of the dummy sex and identity on *approach* and *interaction* separately in *Morpho helenor* and *M. achilles*. The day of experiment was also included in the models to control for stochastic variation during the two-month study. We further tested if variation in wing area and proportion of iridescent blue among dummies affected the frequency of approach and interaction, again using logistic regression analyses (Fig. S7). Statistical significance of each variables was assessed using likelihood ratio tests comparing logistic regression models (Lewis et al., 2011). Finally, we tested the effect of dummy sex and identity on the duration of interaction using Kruskal-Wallis tests.

Based on the flight kinematic data, we investigated whether flight behaviour during the interaction differed with male *vs*. female dummies. We used linear regressions to test how variation in proportion of time spent, speed and acceleration varied with distance from dummy male *vs.* female during the flight interaction. We also tested for differences in mean flight distance from dummy between dummy sex using Wilcoxon test.

## Author Contributions

C.L.R., V.L., and V.D. designed the research. C.L.R., V.L., V.D., C.R., E.A., and H.B. performed the research. C.L.R., V.L., and V.D. wrote the paper.

## Acknowledgments

The authors would like to thank the Peruvian authorities, and in particular SERFOR (the Servicio Nacional Forestal y de Fauna Silvestre) for providing the necessary research permits (permit: 373-2017-SERFOR-DGGSPFFS). We thank Ronald Mori Pezo for help with the mark-recapture experiment. C.L.R. acknowledges financial support by Université de Paris and the Ecole Doctorale FIRE - Program Bettencourt. This work was also supported by a grant from Agence National de la Recherche under the LabEx ANR-10-LABX-0003-BCDiv, in the program “Investissements d’avenir” n ANR-11-IDEX-0004-02” to C.L.R and from the Emergence program of Paris city council to V.L.

---

**Fig. S1.**
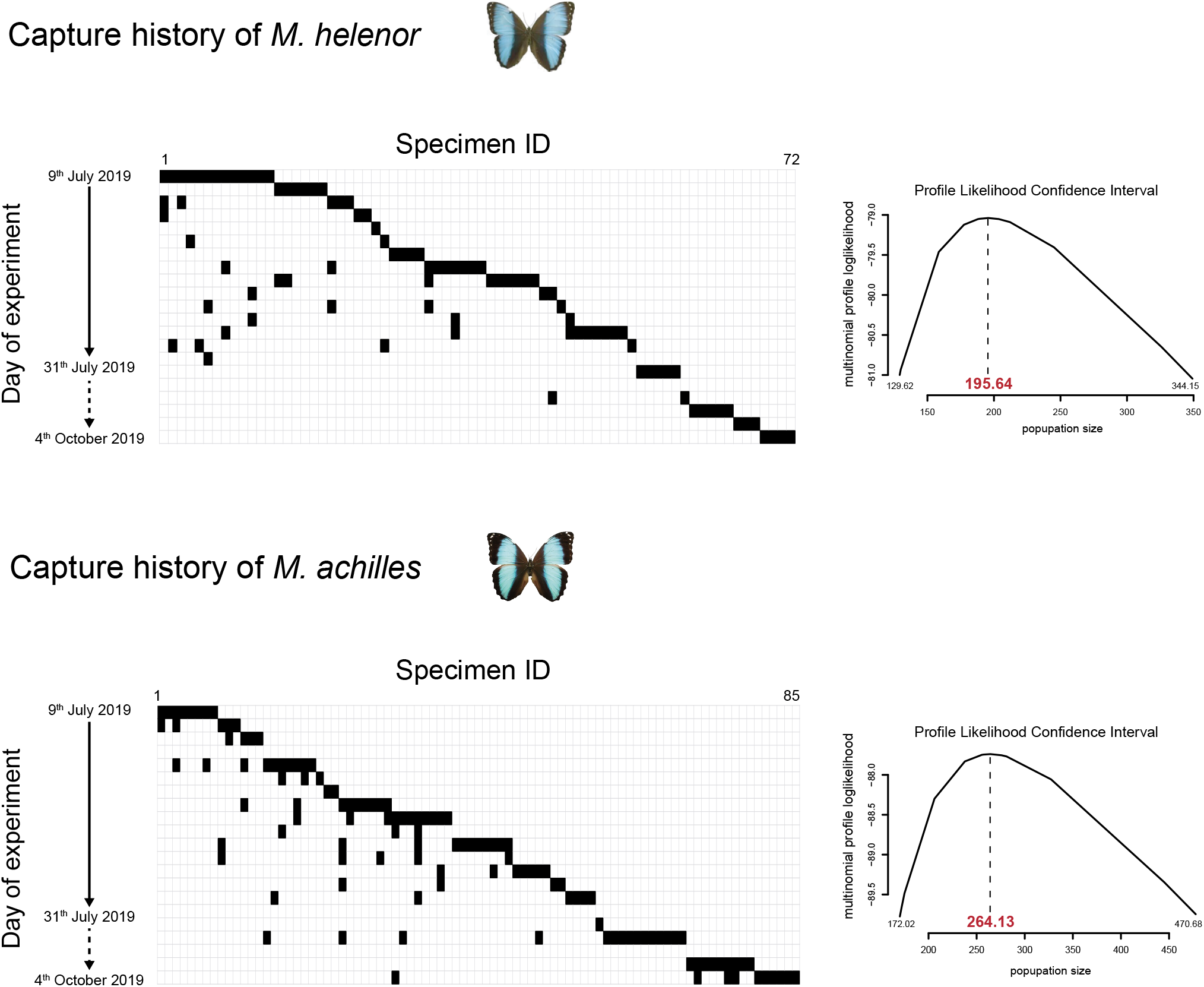
Estimating population size from mark-recapture data. Capture history is shown for the two mimetic sister species *M. helenor* (top) and *M. achilles* (bottom). It gives the capture status on each day of experiment: caught (black boxes) or uncaught (white boxes). Days of experiment along the continuous arrow were nearly consecutive, while those along the dashed arrow were performed every 2 weeks. Based on capture-recapture histories, we estimated individual abundance for each species using a loglinear model implemented in the R package Rcapture (Baillargeon & Rivest 2007), assuming constant population size throughout the experiment. The likelihood confidence interval of the population sizes is shown on the right column.

**Fig. S2.**
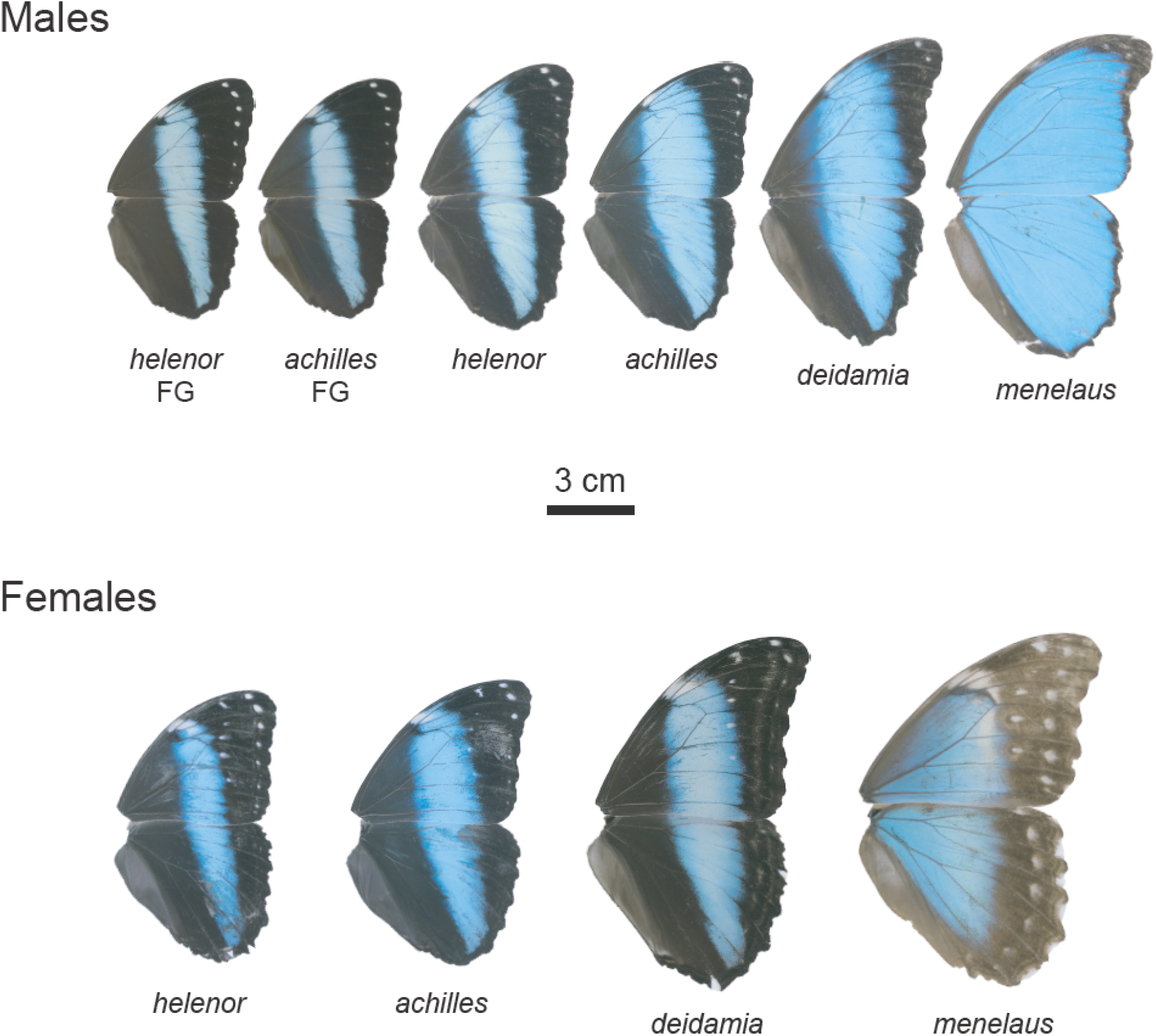
Wings of *Morpho* butterflies used for the dummy experiment. ‘FG’ indicates exotic dummies from French Guiana. All other dummies were build with wings from Peruvian individuals. Wings are shown at their relative size.

**Fig. S3.**
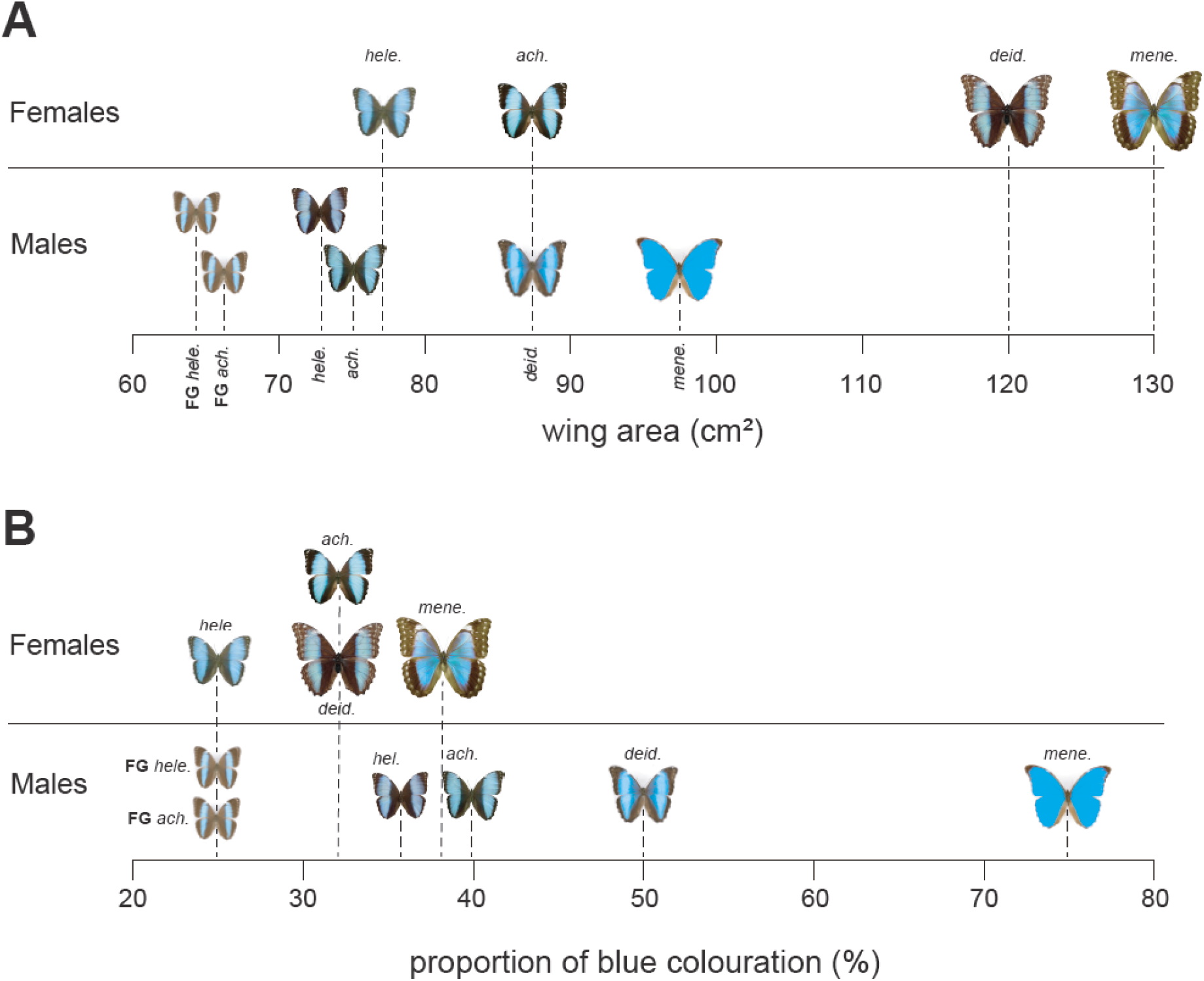
(A) Variation in wing area and (B) in proportion of blue colouration among the tested dummy butterflies. ‘FG’ indicates exotic dummies from French Guiana. All other dummies were build with wings from Peruvian individuals.

**Fig. S4.**
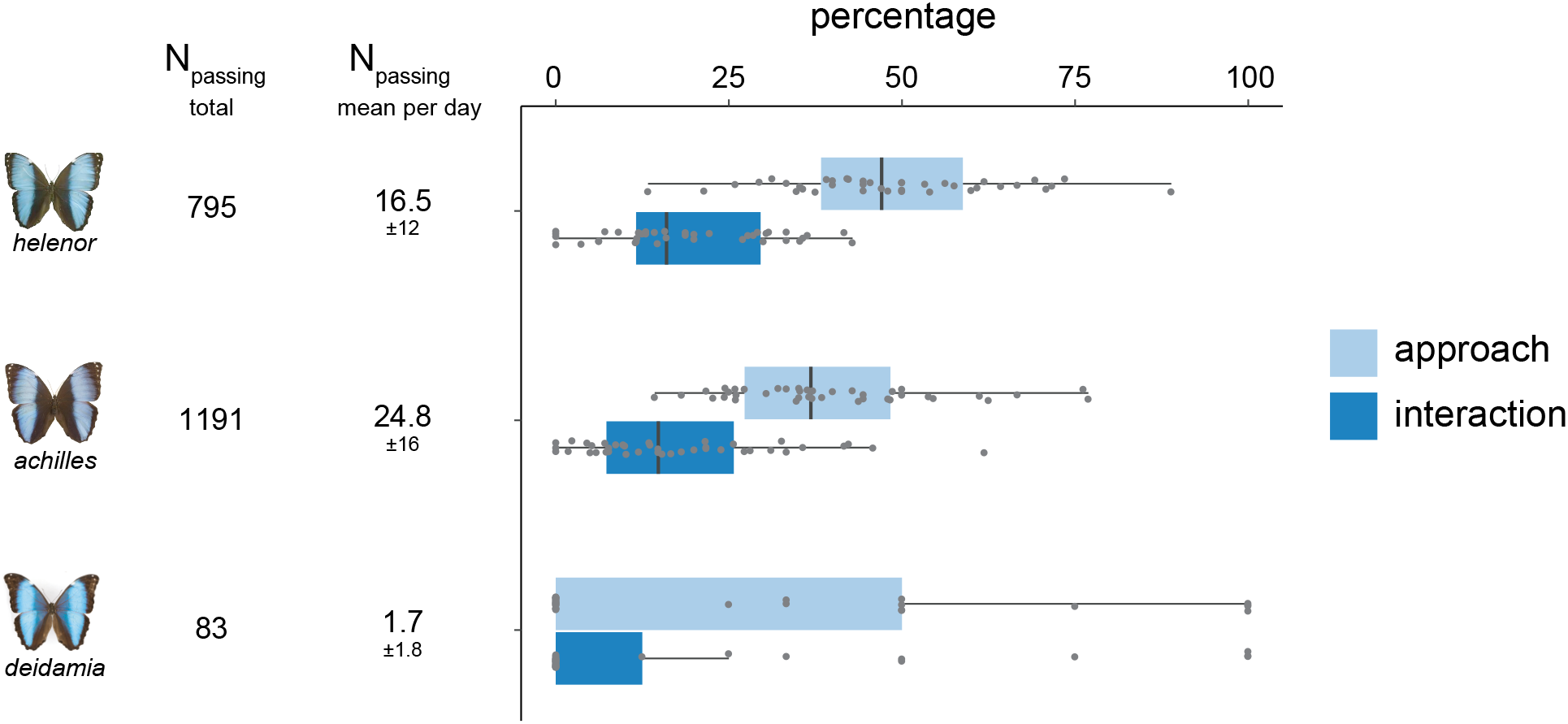
Percentage of approach and interaction with the dummy butterfly (all dummy identity and sex confounded) among sympatric *Morpho* species. Percentage were computed over the number of passing individuals along the river. Total number of passing *Morpho* (in 40 day of experiment) and mean per day is indicated on the left. Each point on the boxplot represent a different day of experiment.

**Fig. S5.**
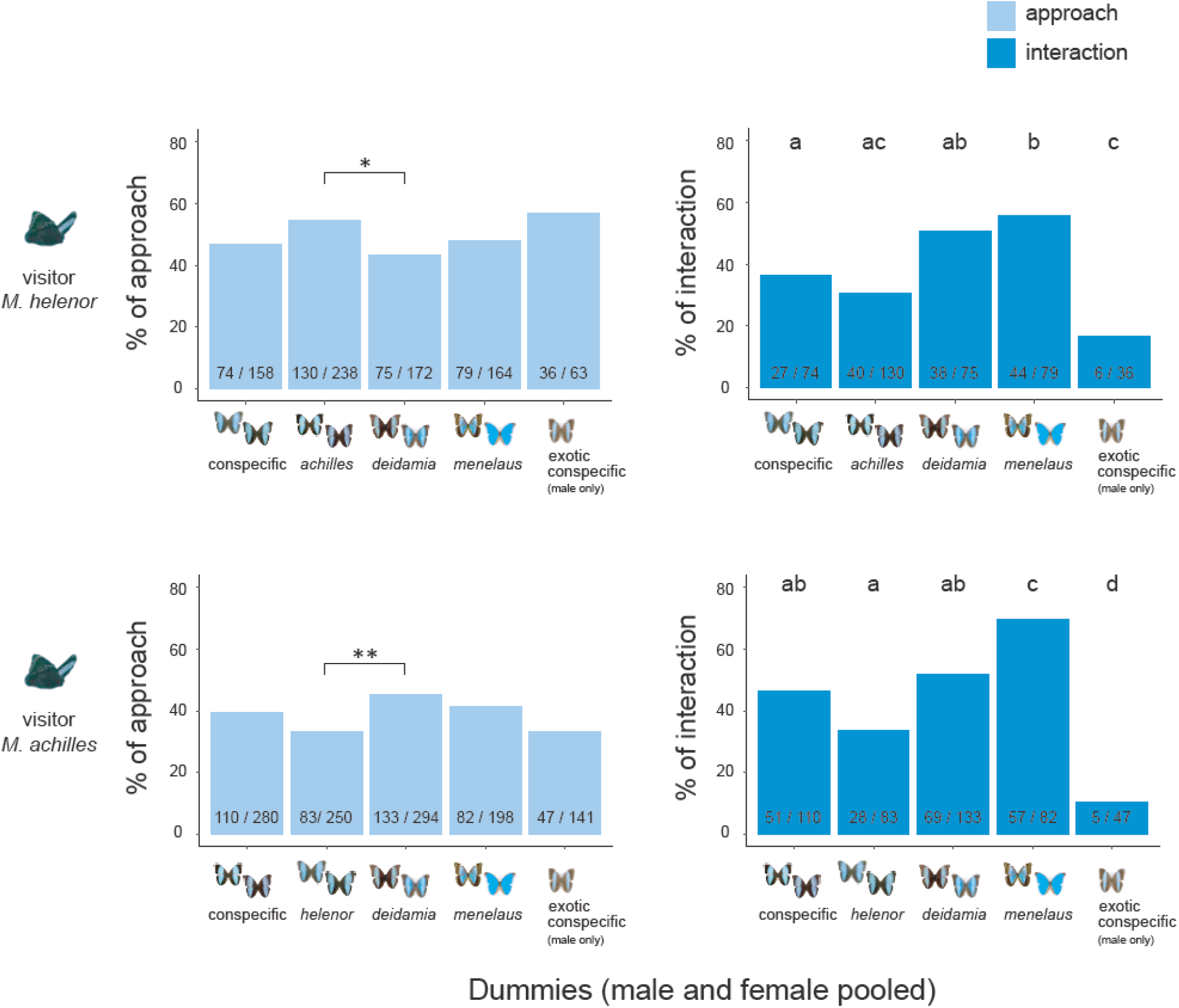
Variation in approach (left column) and interaction (right column) frequency with conspecific and congener dummies in two mimetic sister *Morpho* species (males and females dummies pooled). Female and male dummies are pooled together, excepted for the exotic dummies where only males were tested. Raw data « nb of approaches / nb of passing» are indicated on barplots of the left column. « nb of interaction/ nb of approaches» are indicated on barplot of the right column.

**Fig. S6.**
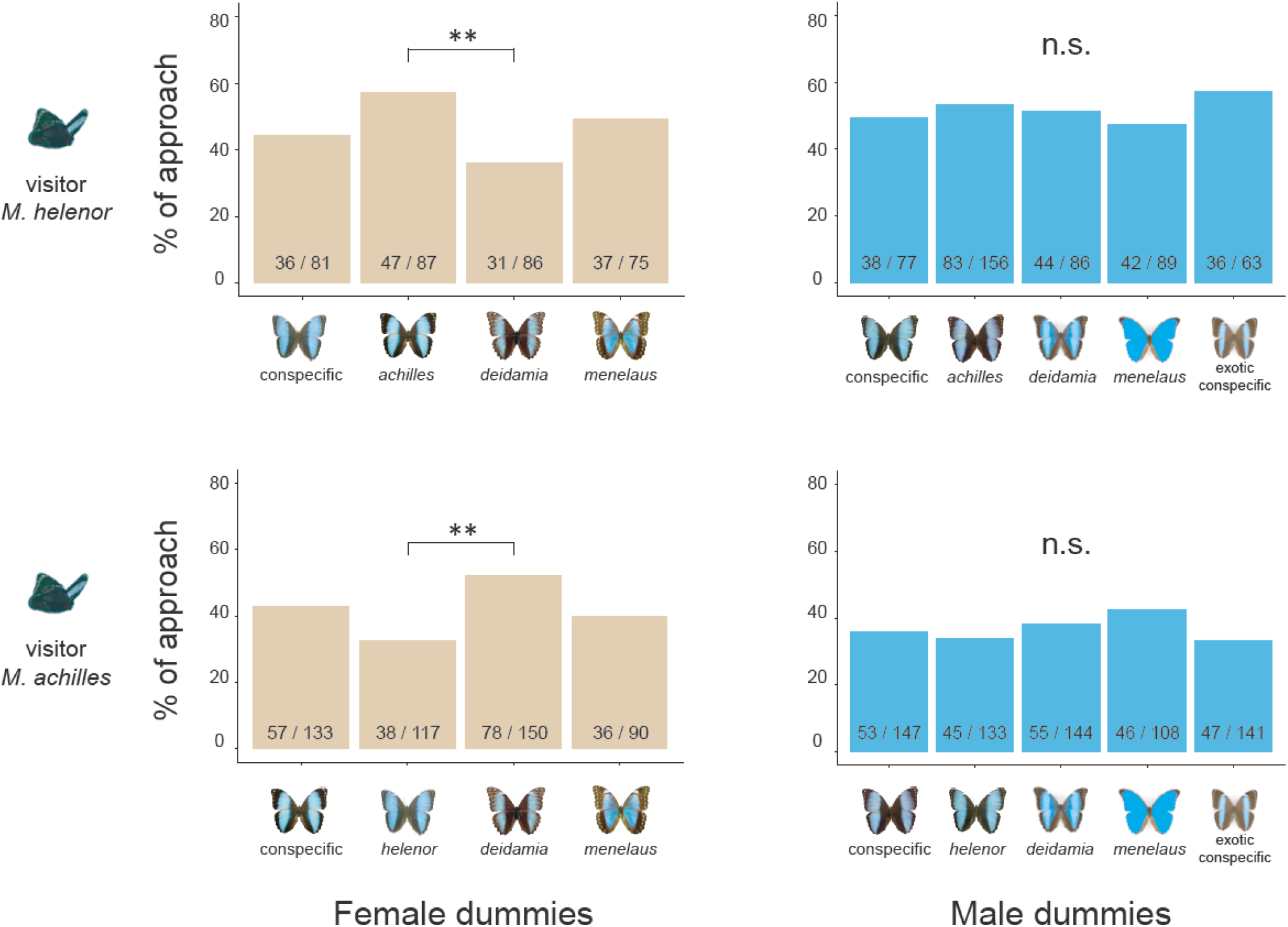
Variation in approach frequency with conspecific and congener dummies in two mimetic sister *Morpho* species. Raw data « nb of approaches / nb of passing » are indicated on each barplot.

**Fig. S7.**
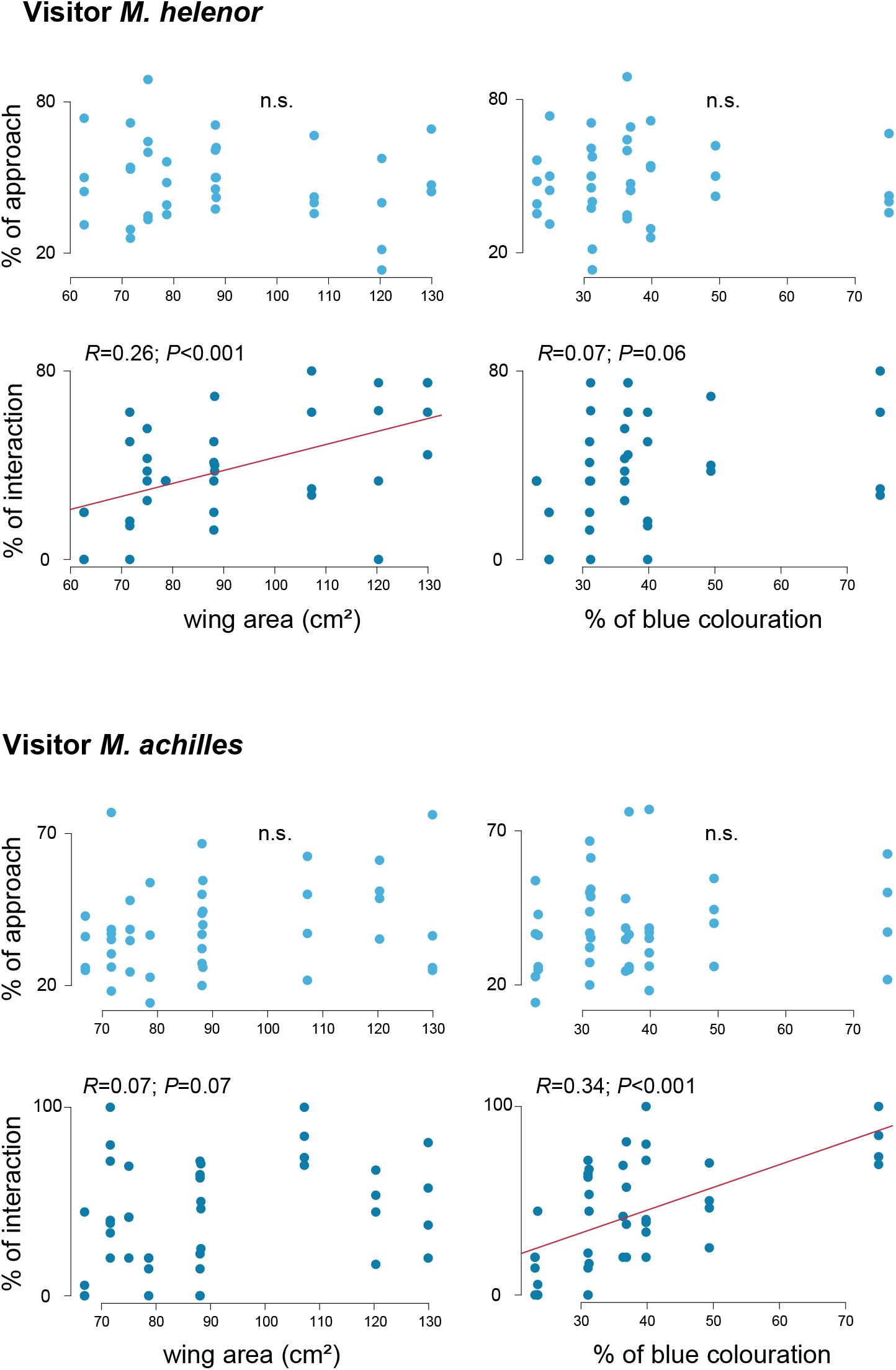
The percentage of approach (top row, light blue) and the percentage of interaction (bottom row, dark blue) are showed in relation to the area and the proportion of blue colouration on the dummy wings. Top panel: visitor *M. helenor.* Bottom panel visitor *M. achilles.* Regression lines are plotted in cases of significant relationships.

**Fig. S8.**
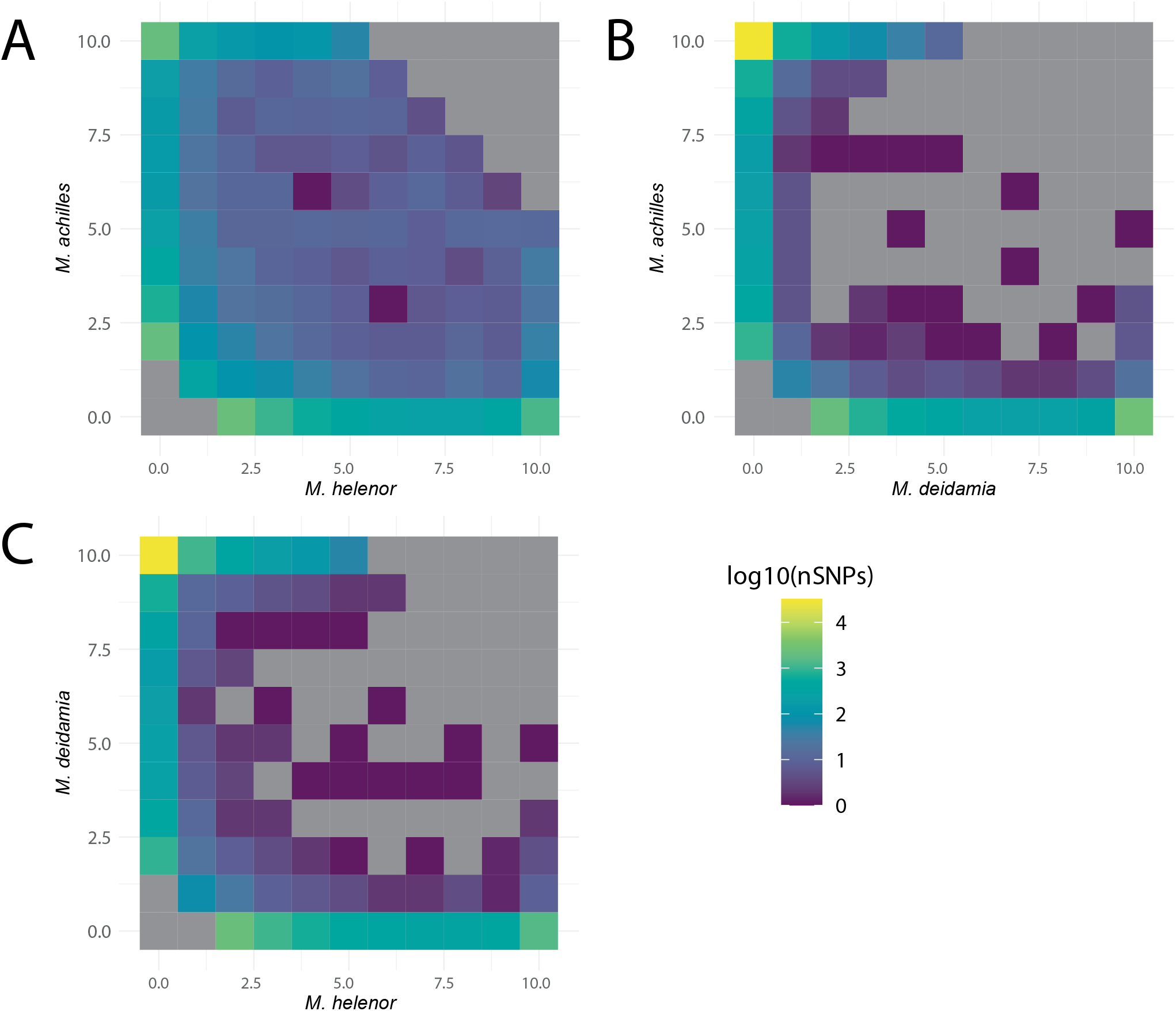
Joint spectra of the allelic frequencies between *M. helenor, M. achilles* and *M. deidamia*. Folded spectrum based on the frequency of the minority allele for each polymorphic position when the three species are aligned. The intraspecific singletons have been removed from the graphical representation to avoid upscaling. Spectra are represented on the log10 scale for three different pairs A) *M. helenor*– *M. achilles*; B) *M. deidamia – M. achilles*; C) *M. helenor – M. deidamia*.

**Fig. S9.**
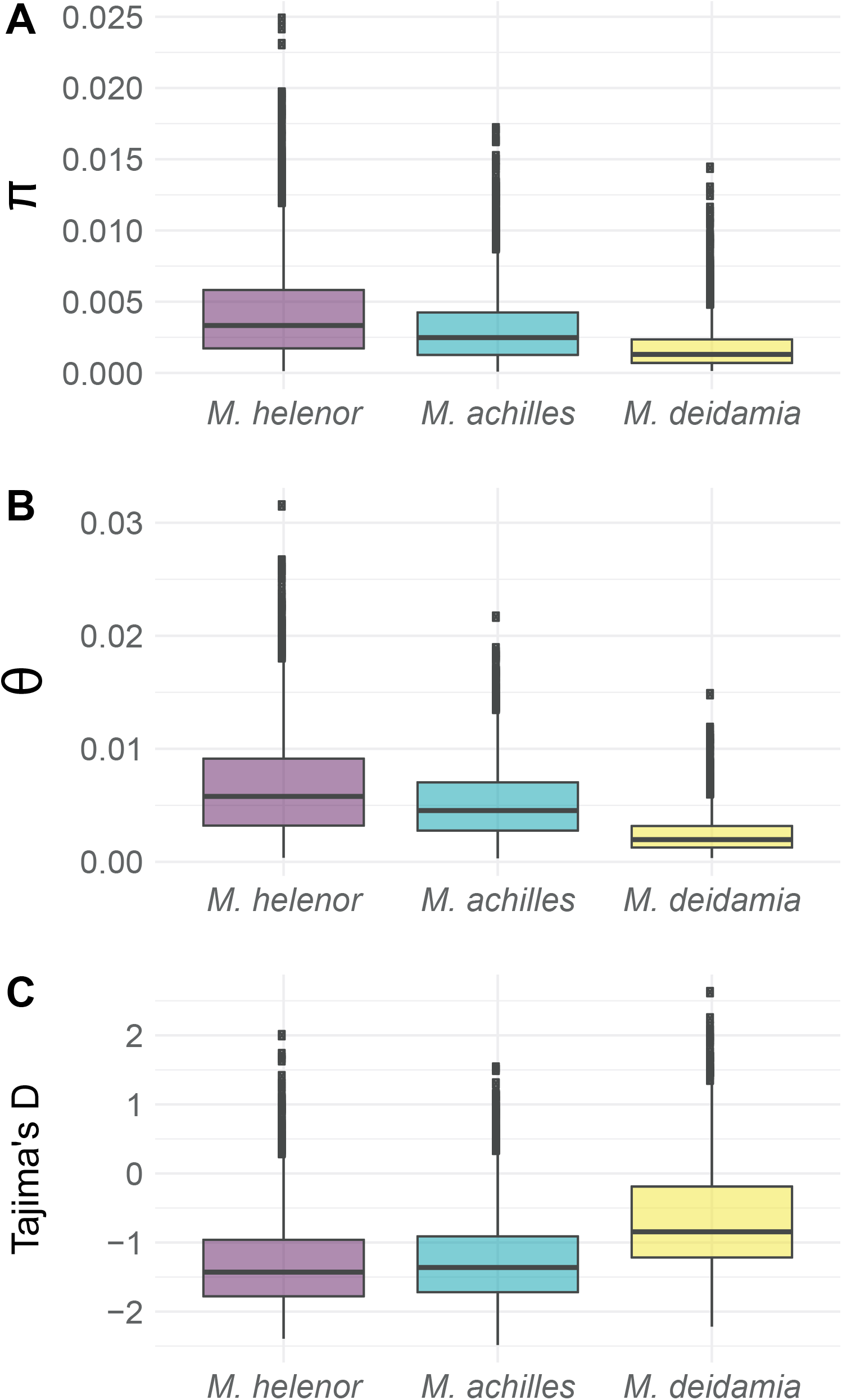
Patterns of within-species molecular diversity. **A**) π (Tajima, 1983); **B**) θ (Watterson, 1975); **C**) Taijma’s D (Tajima, 1989).

**Fig. S10.**
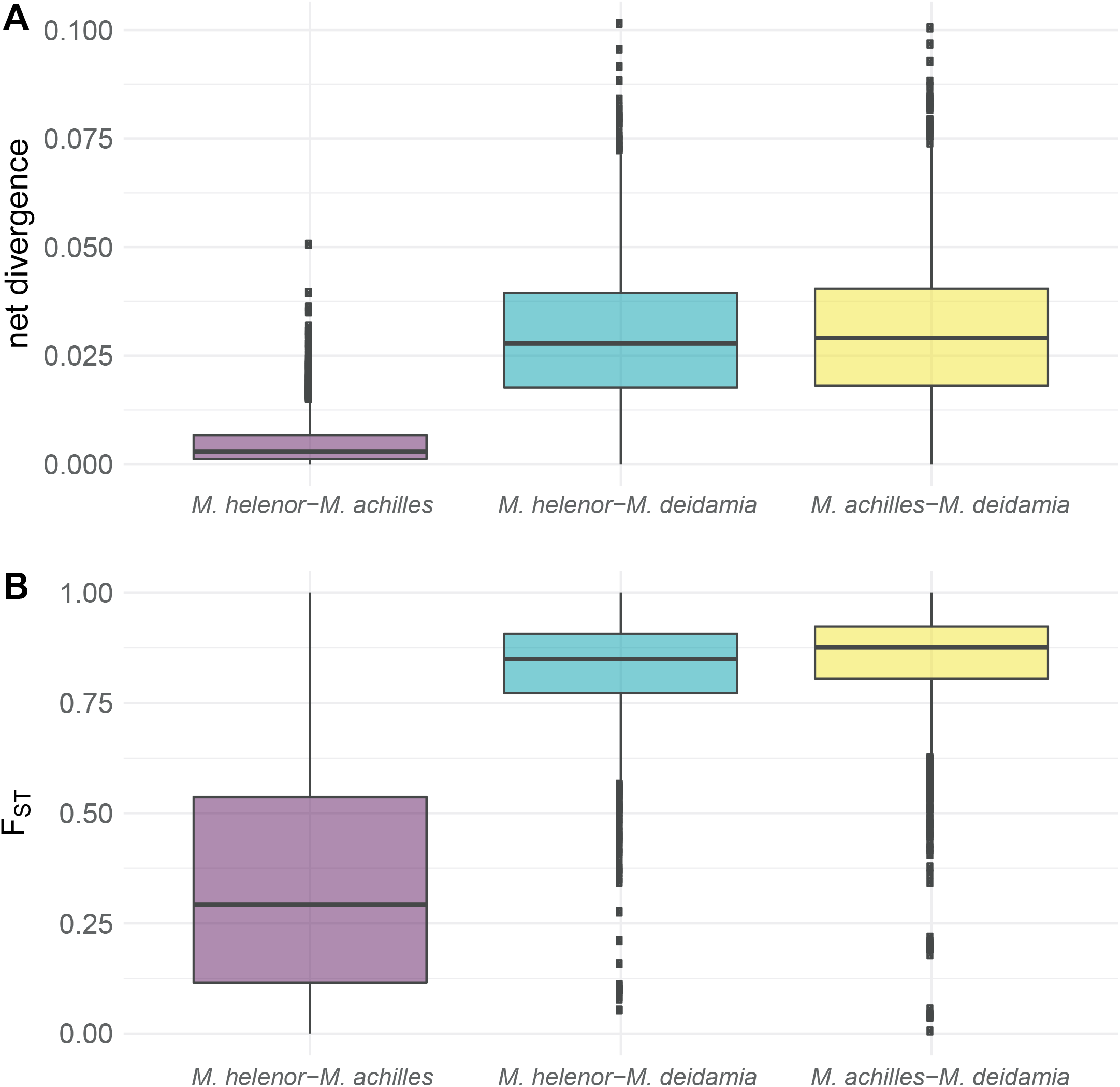
Patterns of between-species divergence and differentiation. A) net divergence measured Da (Nei, 1987). B) Fst computed as (1 - πs) / πT where πs is the average pairwise nucleotide diversity within population and πT is the total pairwise nucleotide diversity of the pooled sample across populations.

**Fig. S11.**
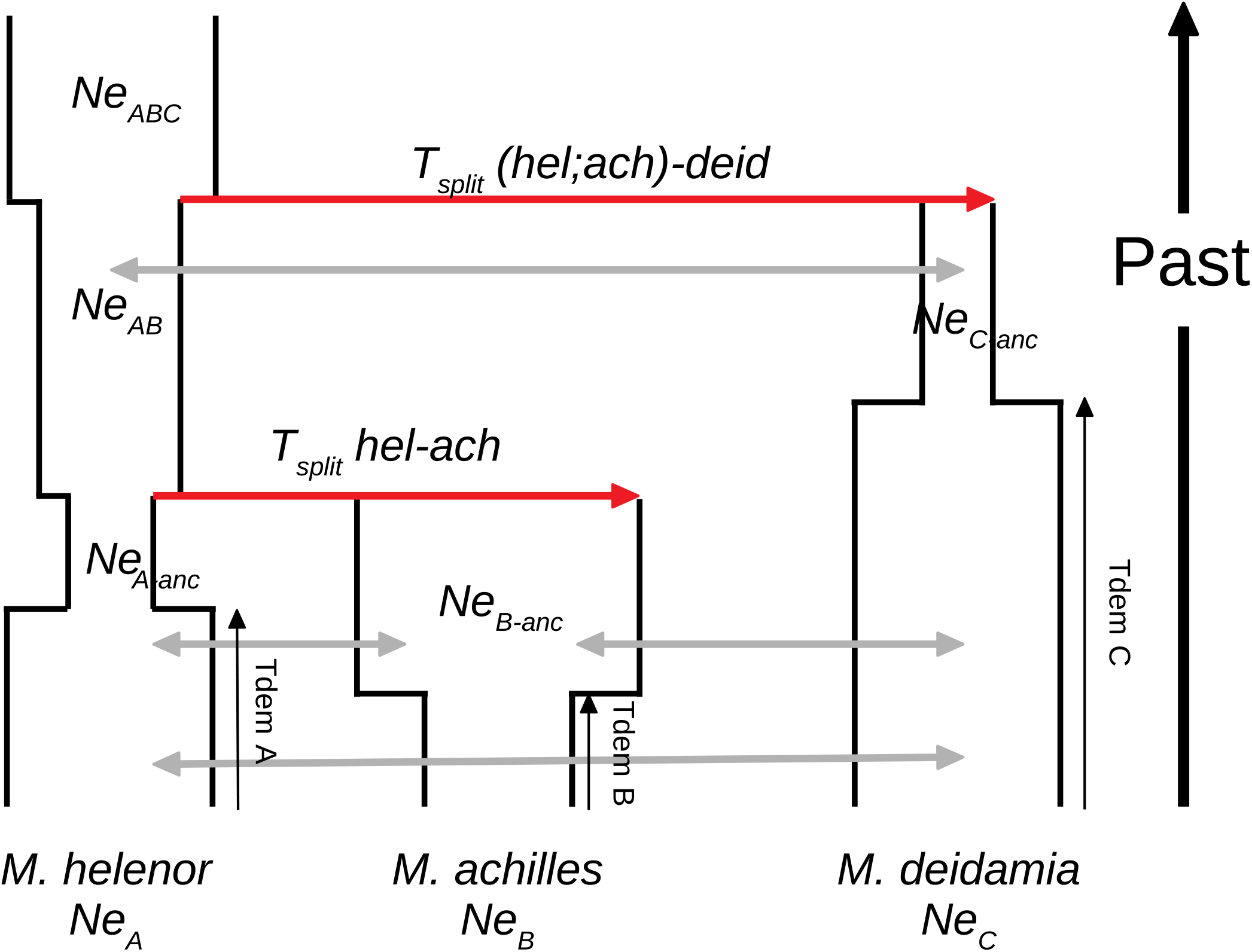
General model of 3 demes speciation with gene flow. This model describes two successive splitting events from the ancestor (size NeABC) to the three sampled species (sizes NeA; NeB; NeC). Each split event is followed by the drawing of a new population size (after the first split: NeAB; NeC–anc. After the second split: NeAC–anc; NeBC–anc). Populations may also undergo a demographic change in size at some point in their recent history, at times Tdem–A, Tdem–B and Tdem–C. This demographic change consists of the sampling of a new population size that may be larger or smaller than the size of their ancestor. Two migration relationships are considered: 1) migration between C and (A;B). 2) migration between A and B. Concerning the first migration relationship, 4 scenarios are explored: migration only between A and C; only between B and C; between A-C and B-C; no migration at all. Concerning the second migration relationship, 2 scenarios are explored: ancient migration (restricted to the first generations after a split) and secondary contact (isolation after a split then contact with gene flow). Ages of migration changes are randomly drawn between zero and the age of the split leading to the concerned lineages.

**Fig. S12.**
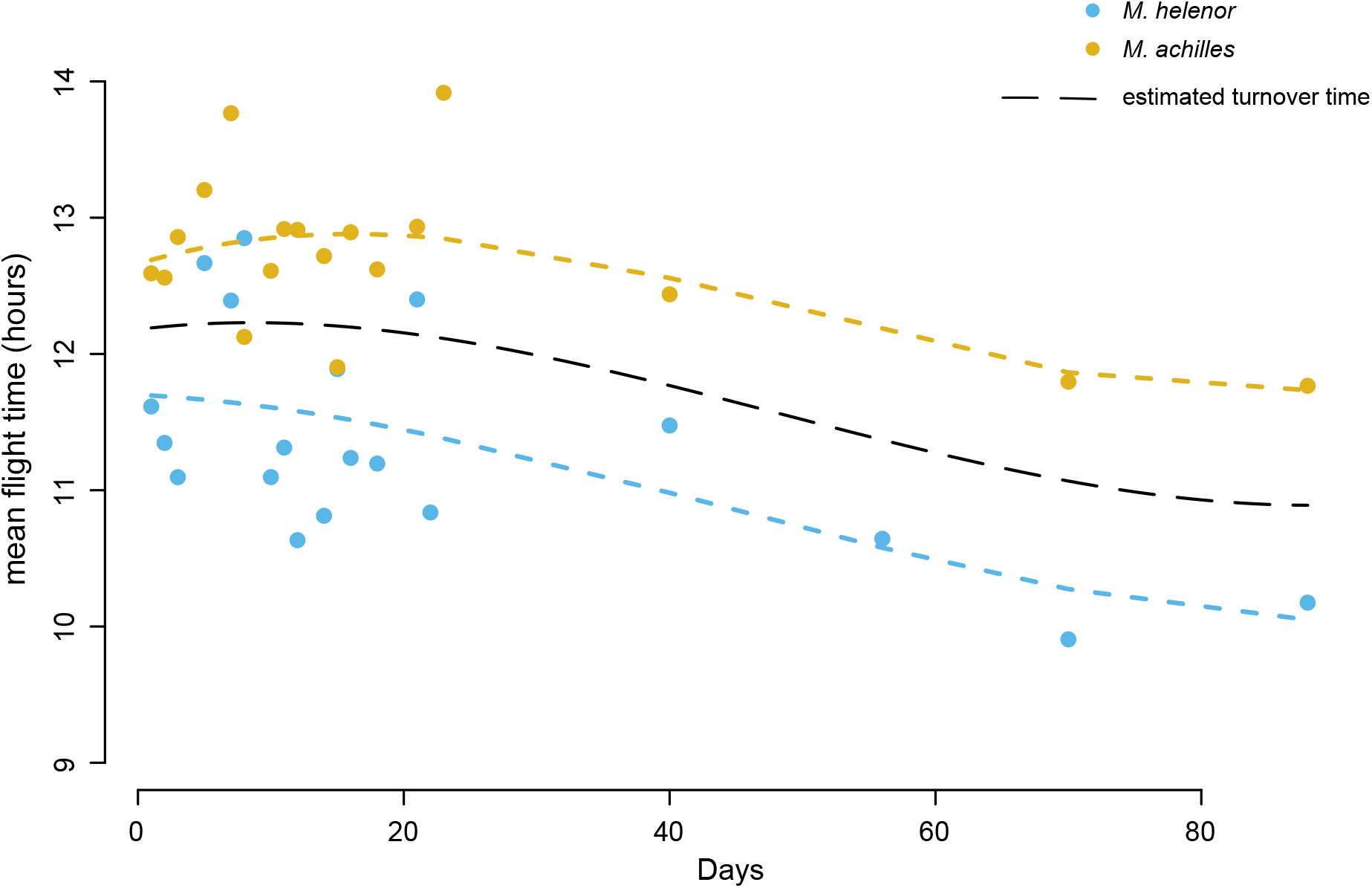
Plot of the mean flight time of the two mimetic species *M. helenor* and *M. achilles* over a ~three-month period. Capture sessions were performed on consecutive days during the first 20 day of experiment. We then performed one day of capture every 2 weeks during 2 months in parallel to the dummy experiment to verify that temporal activity was stable over time. Because turnover time slightly change over the duration of the experiment, we estimated the identify of the flying *Morpho* during the dummy experiment based on the hours of the day and the turnover time estimated for this day.

**Table S1.**
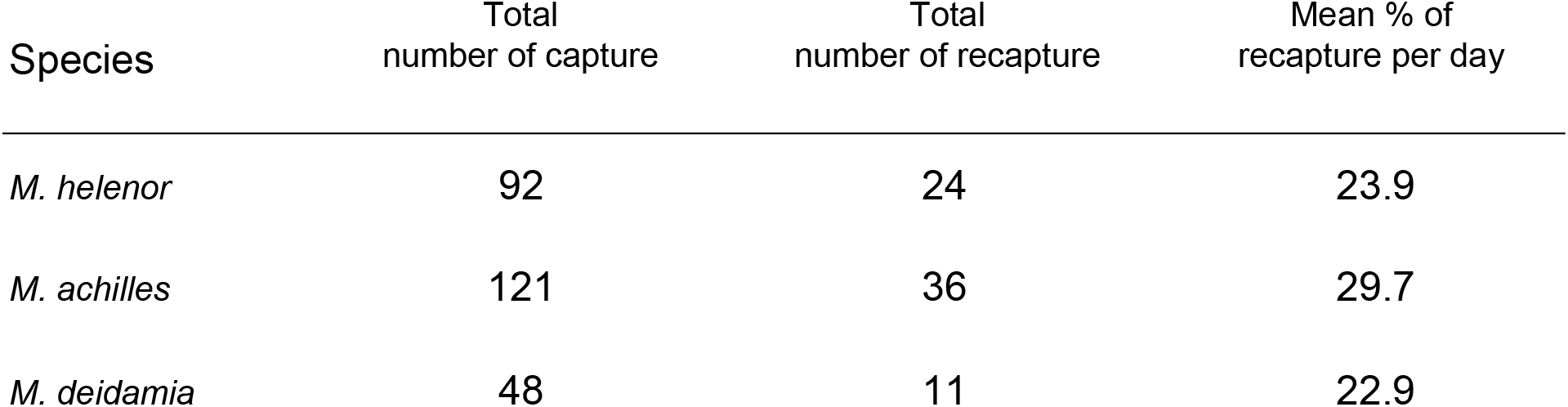
Number of captures and recaptures per *Morpho* species.

| Species | Total<br>number of capture | Total<br>number of recapture | Mean % of<br>recapture per day |
| --- | --- | --- | --- |
| *M. helenor* | 92 | 24 | 23.9 |
| *M. achilles* | 121 | 36 | 29.7 |
| *M. deidamia* | 48 | 11 | 22.9 |

**Table S2.**
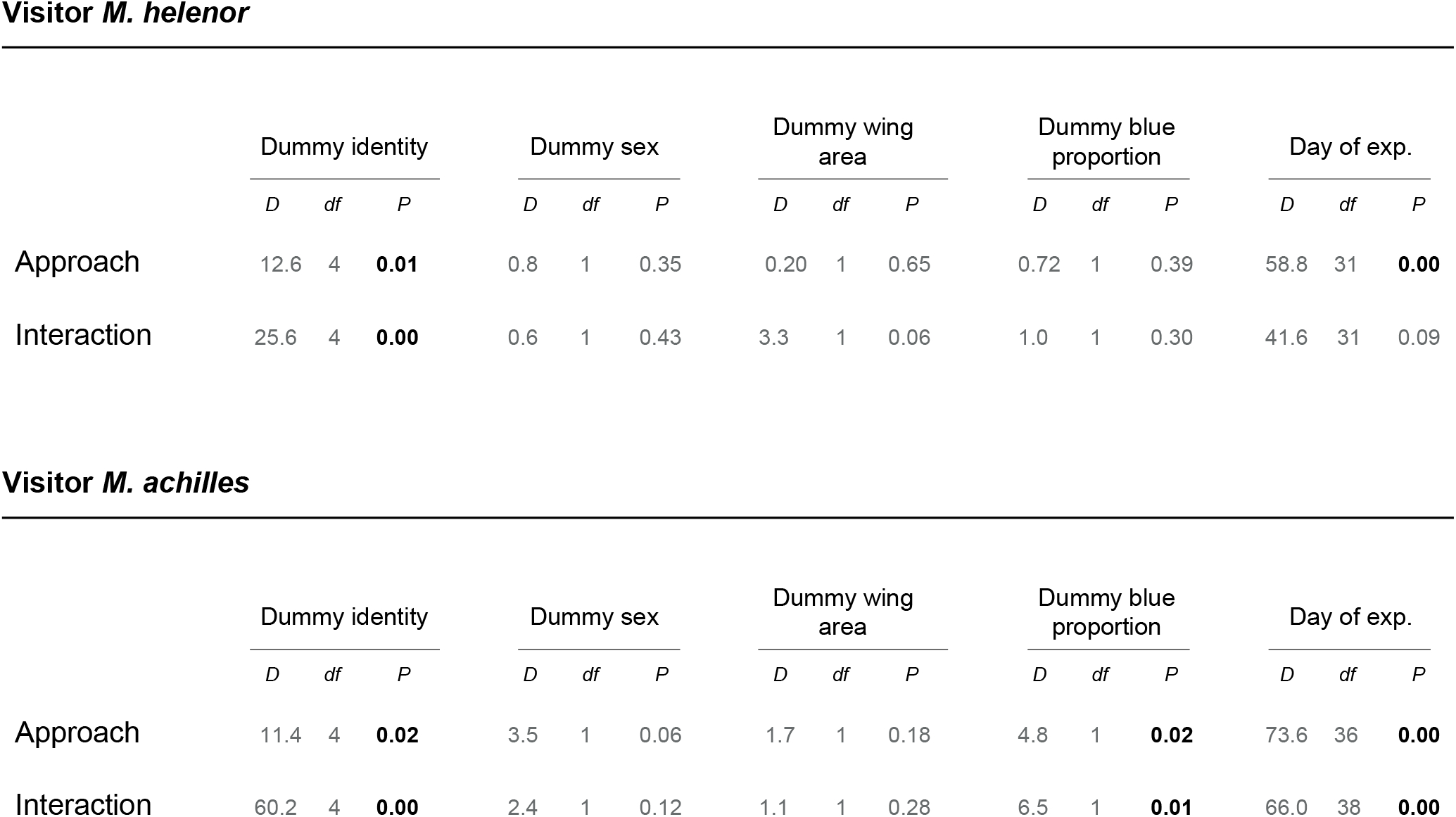
Effect of dummy identity (both sex confounded) on the number of approaches and interactions. The effect of dummy characteristics and of day of experiment on the number of approaches and interactions was tested using logistic regression models. Below are reported the results of likelihood ratio tests comparing models in order to test the global effect of each variable on the number of approaches and interactions.

|  | Dummy identity |  |  | Dummy sex |  |  | Dummy wing area |  |  | Dummy blue proportion |  |  | Day of exp. |  |  |
| --- | --- | --- | --- | --- | --- | --- | --- | --- | --- | --- | --- | --- | --- | --- | --- |
|  | *D* | *df* | *P* | *D* | *df* | *P* | *D* | *df* | *P* | *D* | *df* | *P* | *D* | *df* | *P* |
| Approach | 12.6 | 4 | **0.01** | 0.8 | 1 | 0.35 | 0.20 | 1 | 0.65 | 0.72 | 1 | 0.39 | 58.8 | 31 | **0.00** |
| Interaction | 25.6 | 4 | **0.00** | 0.6 | 1 | 0.43 | 3.3 | 1 | 0.06 | 1.0 | 1 | 0.30 | 41.6 | 31 | 0.09 |

**Visitor *M. achilles***
|  | Dummy identity |  |  | Dummy sex |  |  | Dummy wing area |  |  | Dummy blue proportion |  |  | Day of exp. |  |  |
| --- | --- | --- | --- | --- | --- | --- | --- | --- | --- | --- | --- | --- | --- | --- | --- |
|  | *D* | *df* | *P* | *D* | *df* | *P* | *D* | *df* | *P* | *D* | *df* | *P* | *D* | *df* | *P* |
| Approach | 11.4 | 4 | **0.02** | 3.5 | 1 | 0.06 | 1.7 | 1 | 0.18 | 4.8 | 1 | **0.02** | 73.6 | 36 | **0.00** |
| Interaction | 60.2 | 4 | **0.00** | 2.4 | 1 | 0.12 | 1.1 | 1 | 0.28 | 6.5 | 1 | **0.01** | 66.0 | 38 | **0.00** |

**Table S3.** Effect of dummy identity (sex separated) on the number of approaches and interactions. The effect of dummy identity on the number of approach and interaction was tested using logistic regression models. Below are reported the results of likelihood ratio tests comparing models in order to test the global effect of each variable on the number of approaches and interactions.

|  | Male dummies |  |  |  |  |  | Female dummies |  |  |  |  |  |
| --- | --- | --- | --- | --- | --- | --- | --- | --- | --- | --- | --- | --- |
|  | Dummy identity |  |  | Day of exp. |  |  | Dummy identity |  |  | Day of exp. |  |  |
|  | *D* | *df* | *P* | *D* | *df* | *P* | *D* | *df* | *P* | *D* | *df* | *P* |
| Approach | 4.7 | 4 | 0.31 | 38.1 | 18 | 0.31 | 7.9 | 3 | 0.50 | 19.4 | 12 | 0.07 |
| Interaction | 12.8 | 4 | **0.01** | 32.9 | 18 | **0.01** | 12.1 | 3 | **0.00** | 10.6 | 12 | 0.55 |

**Visitor *M. achilles***
|  | Male dummies |  |  |  |  |  | Female dummies |  |  |  |  |  |
| --- | --- | --- | --- | --- | --- | --- | --- | --- | --- | --- | --- | --- |
|  | Dummy identity |  |  | Day of exp. |  |  | Dummy identity |  |  | Day of exp. |  |  |
|  | *D* | *df* | *P* | *D* | *df* | *P* | *D* | *df* | *P* | *D* | *df* | *P* |
| Approach | 2.7 | 4 | 0.59 | 32.1 | 20 | **0.04** | 14.9 | 3 | **0.00** | 39.9 | 14 | **0.00** |
| Interaction | 55.6 | 4 | **0.00** | 38.1 | 20 | **0.00** | 19.8 | 3 | **0.00** | 28.6 | 14 | **0.01** |

**Table S4.** Effect of wing area and blue proportion of the dummies on the number of approaches and interactions. The effect of dummy wing area and proportion of blue colouration on the number of approaches and interactions was tested using logistic regression models. Below are reported the results of likelihood ratio tests comparing model in order to test the global effect of each variable on the number of approaches and interactions.

**Visitor (all species)**
|  | Dummy wing area |  |  | Dummy blue proportion |  |  | Day of exp. |  |  |
| --- | --- | --- | --- | --- | --- | --- | --- | --- | --- |
|  | *D* | *df* | *P* | *D* | *df* | *P* | *D* | *df* | *P* |
| Approach | 2.60 | 1 | 0.10 | 0.39 | 1 | 0.52 | 132.9 | 46 | **0.00** |
| Interaction | 48.5 | 1 | **0.00** | 12.9 | 1 | **0.00** | 117.1 | 46 | **0.00** |

**Visitor *M. helenor***
|  | Dummy wing area |  |  | Dummy blue proportion |  |  | Day of exp. |  |  |
| --- | --- | --- | --- | --- | --- | --- | --- | --- | --- |
|  | *D* | *df* | *P* | *D* | *df* | *P* | *D* | *df* | *P* |
| Approach | 1.7 | 1 | 0.18 | 0.98 | 1 | 0.33 | 70.5 | 36 | **0.00** |
| Interaction | 22.1 | 1 | **0.00** | 0.12 | 1 | 0.72 | 47.1 | 36 | 0.10 |

**Visitor *M. achilles***
|  | Dummy wing area |  |  | Dummy blue proportion |  |  | Day of exp. |  |  |
| --- | --- | --- | --- | --- | --- | --- | --- | --- | --- |
|  | *D* | *df* | *P* | *D* | *df* | *P* | *D* | *df* | *P* |
| Approach | 11.0 | 1 | **0.00** | 0.0 | 1 | 0.82 | 83.1 | 40 | **0.00** |
| Interaction | 28.3 | 1 | **0.00** | 30.1 | 1 | **0.00** | 83.9 | 40 | **0.00** |

**Table S5.**
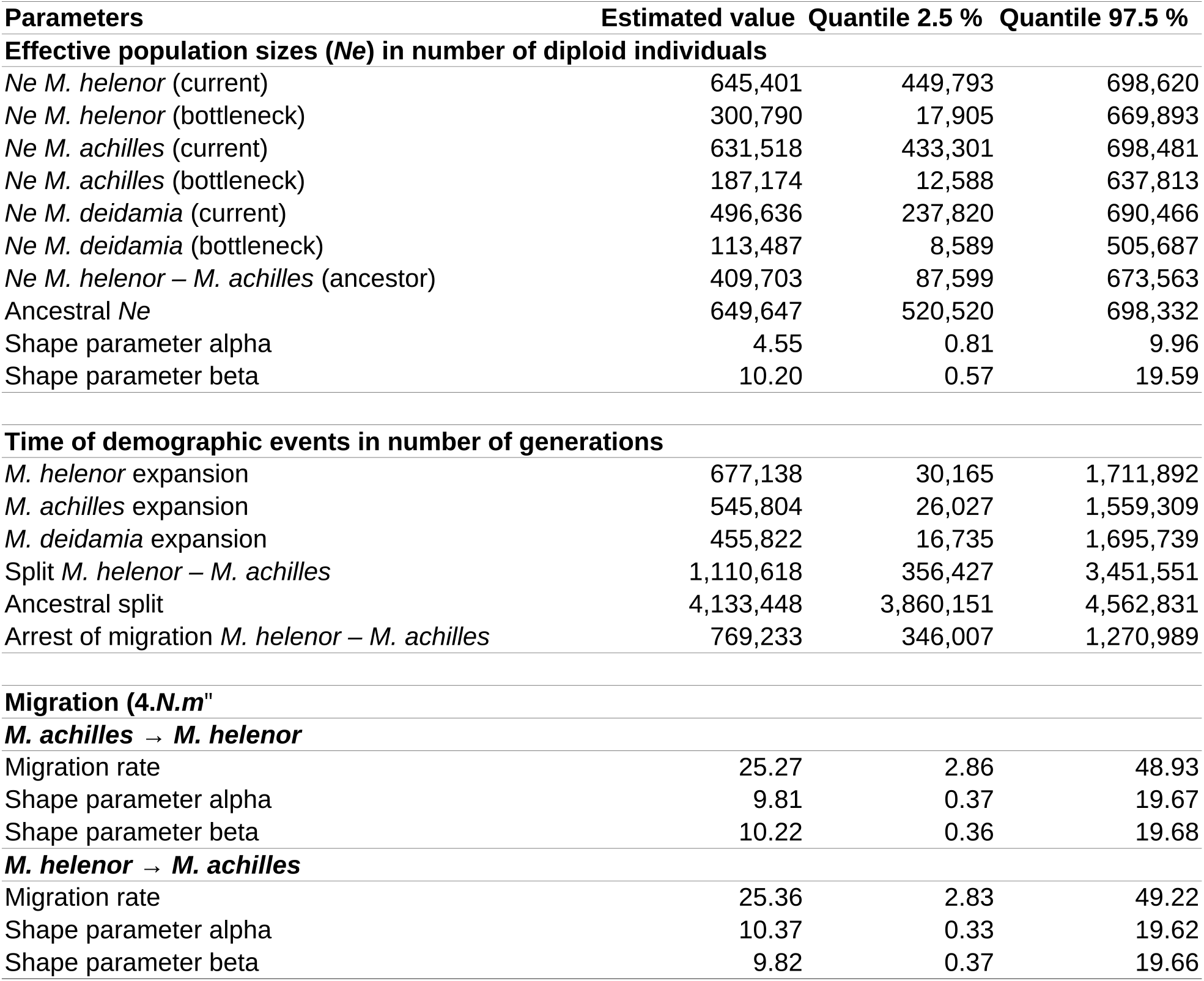
Parameters inferred by Random Forest for the best fitting model (Fig. 4; (Raynal et al., 2019)).

